# The intrinsically disordered N-terminus of SUMO1 is an intramolecular inhibitor of SUMO1 interactions

**DOI:** 10.1101/2024.01.04.574143

**Authors:** Sebastian M. Richter, Fan Jin, Tobias Ritterhoff, Aleksandra Fergin, Eric Maurer, Andrea Frank, Michael Daube, Alex Hajnal, Rachel Klevit, Frauke Gräter, Annette Flotho, Frauke Melchior

## Abstract

Ubiquitin-related proteins of the SUMO family are reversibly attached to thousands of proteins in eukaryotic cells. Many SUMO substrates, effectors and enzymes carry short motifs (SIMs) that mediate low affinity interactions with SUMO proteins. How specificity is achieved in target selection, SUMO paralogue choice and SUMO-dependent interactions is largely unknown. A unique but poorly understood feature of SUMO proteins is their intrinsically disordered N-terminus. We reveal a function for N-termini of human, *C. elegans,* and yeast SUMO proteins as intramolecular inhibitors of SUMO-SIM interactions. Mutational analyses, NMR spectroscopy, and Molecular Dynamics simulations indicate that SUMO’s N-terminus can inhibit SIM binding by fast and fuzzy interactions with SUMO‘s core. Deletion of the *C. elegans* SUMO1 N-terminus leads to p53-dependent apoptosis during germline development, indicating an important role in DNA damage repair. Our findings reveal a mechanism of disorder-based autoinhibition that contributes to the specificity of SUMOylation and SUMO-dependent interactions.

## INTRODUCTION

SUMOylation, the reversible attachment of small ubiquitin-related modifier SUMO to target proteins, serves as a molecular switch that regulates protein functions. It is particularly important in pathways that contribute to chromatin organization and function including transcription, replication, DNA damage repair, nucleocytoplasmic transport, and chromosome segregation^1–4^. At least a thousand proteins are subject to this essential protein modification under physiological conditions, and many more in severe stress (e.g., ^5,6^). The mechanism of SUMOylation resembles that of Ubiquitylation: Isopeptide bond formation between the C-terminal di-glycine motif of SUMO and the ε-amino group of a lysine sidechain in target proteins is catalyzed by an enzymatic trio of the SUMO-specific E1 activating heterodimer Aos1/Uba2 (SAE1/SAE2), the SUMO E2 conjugating enzyme Ubc9, and usually one of several SUMO E3 ligases. SUMO-specific isopeptidases reverse the modification. One striking difference to the Ubiquitin system is the small number of enzymes involved. Over 600 E3 ligases are known for Ubiquitylation but fewer than ten have been identified for SUMOylation, amongst them the Siz/PIAS RING E3 ligases, ZNF451, and the nucleoporin RanBP2^1,7^. Similarly, nearly one hundred isopeptidases with specificity for Ubiquitin are known, yet only eight enzymes appear to act on mammalian SUMO^8,9^. On the other hand, many organisms have more than one SUMO protein: Whereas *S. cerevisiae, D. melanogaster, and C. elegans* harbor only one SUMO gene, the weed *A. thaliana* expresses 8 SUMO genes (reviewed in ^10^) and the human genome encodes five SUMO paralogues, three of which are ubiquitously expressed, SUMO1 and the twins SUMO2 and SUMO3 (reviewed in ^7^). SUMO2 and 3 are virtually identical but share only about 50% identity with SUMO1. The two classes of SUMO proteins differ in expression levels, response to cellular stress, and their ability to form SUMO chains. The N-termini of human SUMO2 and 3 as well as *S. cerevisiae* Smt3 contain at least one lysine residue, on which SUMO chains can assemble^11–13^, whereas the human SUMO1 and *C. elegans* SMO-1 N-termini are devoid of lysine (see also below). Although many proteins can be modified *in vitro* and in cells with both classes of SUMO proteins, a significant number of proteins is preferentially or exclusively modified with one or the other SUMO paralogue at endogenous expression levels in cells (e.g. ^14^).

How paralogue-specific modification and effector binding are brought about is only partially understood. Both events require SUMO-binding (reviewed, e.g. in ^7^). A SUMO-binding motif in the target may help to recruit the Ubc9 - SUMO thioester. If this motif prefers a specific SUMO paralogue, the target is also preferentially modified with this paralogue as in USP25 and BLM^15,16^. SUMO-binding motifs in downstream effectors help to recruit specific SUMOylated proteins. Examples are the recruitment of Srs2 to SUMOylated PCNA^17^ and the recruitment of the ubiquitin ligase RNF4 to polySUMOylated PML^18,19^.

To date, at least three different classes of SUMO interactions have been identified, of which the best studied class I interactions involve SUMO Interaction Motifs (SIMs)^20^.

Known examples for class II interactions with a surface of SUMO opposite of the SIM binding site are Dpp9 and Ubc9 ^7,21–24^, while ZZ domains have been shown to either bind SUMO’s core as in the case of CBP^25^ or the absolute N-terminus as for HERC2^26^. SIMs typically consist of a four amino acid patch with three hydrophobic residues, most commonly valine or isoleucine, and a variable at position 3 ([P/I/L/V/M]-[I/L/V/M]- x- [I/L/V/M]), followed by a stretch of negatively-charged amino acids and/or serine residues^27^. In some cases, the orientation of the SIM can be reversed. A distinct subclass of SIMs is characterized by a [V/I/L/F/Y]-[I/V]-D-L-T consensus motif^28^. The hydrophobic amino acids in the core of SIMs insert as a β-strand into the hydrophobic groove formed by the β2-strand and the α1-helix of SUMO (SIM-binding groove)^29,30^. An exception is TDP2, which binds the SIM-binding area of SUMO2/3 via five separate binding elements, two β-sheets and three surface loops, forming a so-called “split SIM”^31^. As is common for binding modules that recognize Ubiquitin or Ubls^32^, SUMO - SIM interactions are usually of low affinity (KD between 1 - 100 μM)^20^. Biological relevance comes about by combining multiple SIMs (as in RNF4), combining a SIM with a second interaction module specific for a given substrate (as is true for Srs2, which recognizes both SUMO and PCNA), or through posttranslational modifications in proximity to the SIM that increase the affinity for SUMO (e.g., phosphorylation in proximity to a SIM in PIAS1^33^). How paralogue-specific interactions are established in SUMO - SIM interactions remains enigmatic. Most of the amino acids in the SIM-binding groove that contribute to SUMO - SIM interactions by NMR are conserved between SUMO1 and SUMO2/3^34–38^. Initially it was proposed that a patch of negative amino acids adjacent to the SIM is required for SUMO1 - but dispensable for SUMO2/3 - interaction^29^. However, USP25^15^ and ATF7IP^38^ exhibit a high preference for SUMO2/3 despite a patch of negative amino acids adjacent to their SIM. Attempts to convert SUMO2/3 into a SUMO1 interacting protein by mutagenesis have thus far been unsuccessful (e.g., ^38^). A defining feature of SUMO proteins is the intrinsically disordered N-terminus^39^, whose function is only partly understood^7,40–42^. While N-termini of the SUMO2/3 family contain a SUMOylation consensus motif (ΨKxE, where Ψ is a bulky hydrophobic amino acid) used for SUMO chain formation^11–13^, this motif is lacking in SUMO1 proteins. In yeast, polymeric SUMO chains contribute to chromatin regulation^40,43^ but the N-terminus itself is dispensible for essential SUMO functions^41^. Here, we reveal an unexpected function for the intrinsically disordered N-termini as intramolecular inhibitors of SUMO - SIM interactions.

## RESULTS

### Intrinsically disordered N-termini of SUMO1 proteins inhibit protein interactions

All SUMO orthologs from yeast to human carry an intrinsically disordered N-terminal extension^39^, typically in the range of 13 - 23 amino acids, as shown in Figure 1A for SUMO1 (adapted from ^39^) and as indicated by boxed residues in the primary structures of the four SUMO proteins used in this study, human SUMO1, human SUMO2, *C. elegans* SMO-1, and *S. cerevisiae* Smt3 (Fig. 1B). Previously, we had performed pull-down screens that allowed us to identify paralogue - specific SUMO- binding partners^15^. In these experiments, we also included a SUMO1 variant lacking the first 19 amino acids as we hypothesized that the N-terminus contributes to interactions. To our surprise, the contrary was the case - the pulldown with SUMO1ΔN19 yielded more proteins than the pull-down with SUMO1 (Fig. S1A + S1B). To test whether SUMO’s disordered N-terminus indeed prevents protein interactions, we conducted pull-down assays with validated binding partners and wild-type SUMO1, the N-terminally truncated variants SUMO1ΔN19 and SUMO1ΔN10, wild-type SUMO2, and SUMO2ΔN13 (Fig. 1C - F). All SUMO variants were untagged and immobilized on CNBr-activated Sepharose in random orientation. Binding partners included two proteins with preference for SUMO2/3, the canonical SIM-containing protein USP25^15^, and TDP2, which has an unusual "split SIM"^31^, the SUMO1-specific SIM-containing RanBP2 fragment RanBP2ΔFG^44,45^ and Ubc9, which binds SUMO1 and SUMO2/3 via a region distinct from the SIM binding groove^46^. Indeed, deletion of the complete SUMO1 N-terminus dramatically increased interaction with USP25 (Fig. 1C + D). Similarly, interaction with TDP2 was enhanced when SUMO1‘s N-terminus was removed, while the interaction with Ubc9 was unaffected. Some increase was observed when half the SUMO1 N-terminus was deleted. As shown in Figure 1E and 1F, deletion of SUMO1‘s N-terminus allowed USP25 to bind SUMO1 as efficiently as wt SUMO2. Importantly, deletion of SUMO2‘s N-terminus (ΔN13) did not increase but not SUMO2‘s - N-terminus inhibits the interaction. To exclude artefacts due to the SUMO immobilization, we repeated pulldown experiments with C-terminally GST-tagged SUMO variants immobilized on Glutathione beads (Fig. S1C + S1D). Again, USP25 bound to ΔN19 SUMO1 as efficiently as to SUMO2, and much better than to SUMO1.

**Figure 1.**
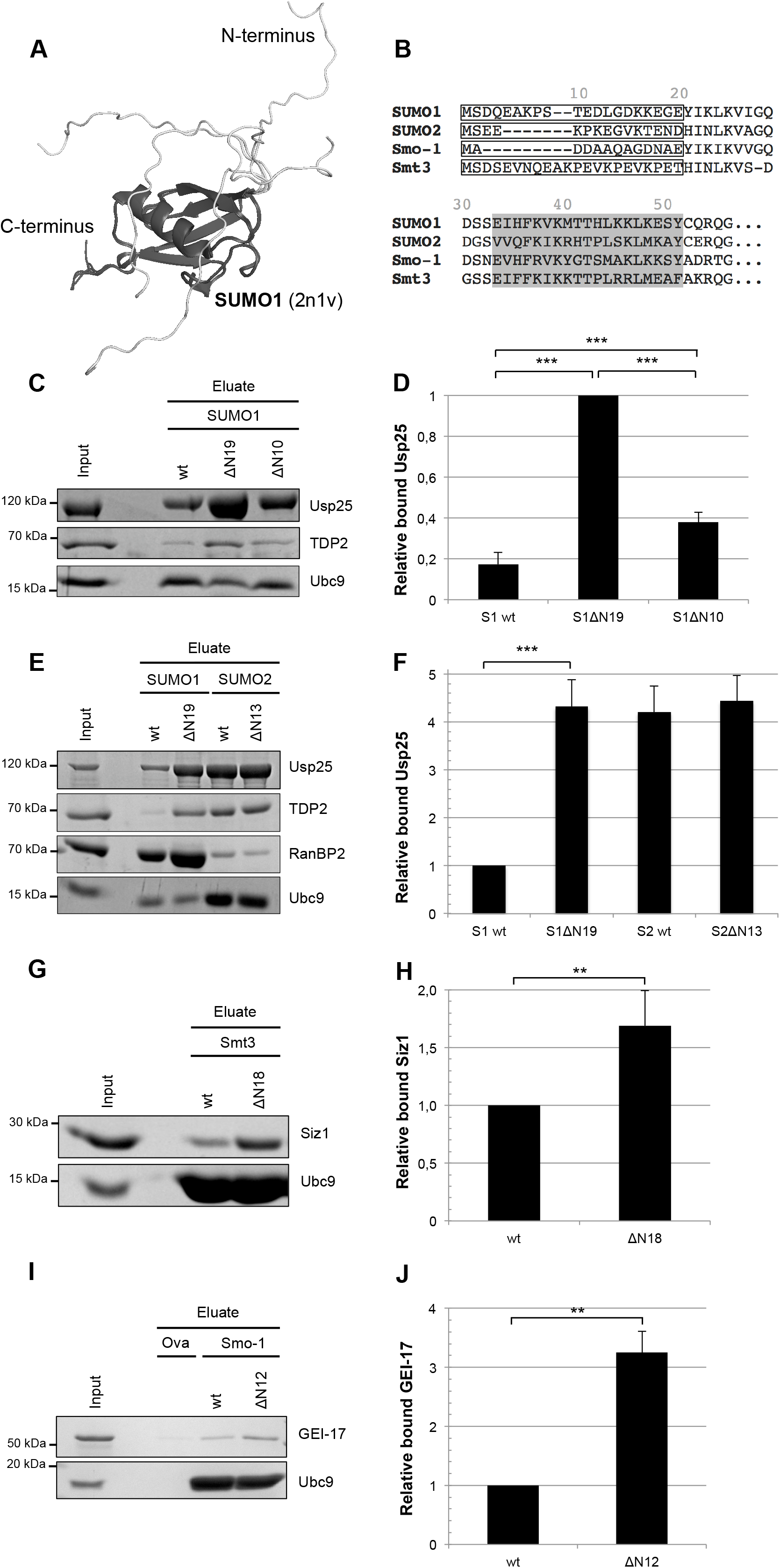
N-terminal deletions of SUMO1 proteins, but not SUMO2, greatly increase binding of SIM-containing proteins. (A) Overlay of 5 selected states of a SUMO1 solution structure (2n1v) using Pymol. Whereas the SUMO core and C-terminus (dark grey) remain stable in all states, the N- terminus (light grey) shows a high degree of flexibility. (B) Partial sequence alignment of human SUMO1, SUMO2, *C. elegans* SMO-1, and *S. cerevisiae* Smt3 using the Clustal Omega web tool. The flexible N-terminal extension is indicated by a box, the SIM-binding groove is highlighted grey. (C) and (E) Pulldown of recombinant Usp25, Ubc9, TDP2 or RanBP2 with SUMO1 or SUMO2 analyzed by SDS-PAGE and Coomassie staining. (D) The amounts of Usp25 in the eluates of SUMO pull-downs in (C) were quantified relative to Ubc9 (n = 4). (F) The amounts of Usp25 in the eluates of SUMO pull-downs in (E) were quantified relative to Ubc9 (n = 6). (G) Pulldown of recombinant *S. cerevisiae* Siz1 or hUbc9 with Smt3 analyzed by SDS-PAGE and Coomassie staining. (H) The amounts of Siz1 were quantified relative to hUbc9 (n = 4). (I) Pulldown of recombinant *C. elegans* GEI-17 or hUbc9 with Ovalbumin (Ova) or Smo- 1 beads analyzed by SDS-PAGE and Coomassie staining. (J) The amounts of GEI-17 were quantified relative to hUbc9 (n = 3). Error bars represent one standard deviation. Asterisks indicate p-values: n.s.: P > 0.05; **: P ≤ 0.01; ***: P ≤ 0.001

To address whether the inhibitory role of the unstructured N-terminus is a unique feature of mammalian SUMO1 or whether it may be conserved in evolution, we turned to yeast and *C. elegans*. The single yeast SUMO protein Smt3 shares about 45% sequence identity with both SUMO1 and SUMO2/3. While hsSUMO1 can complement loss of Smt3, hsSUMO2 and 3 fail to do so, due to incompatibility of these human SUMO proteins with some of the yeast SUMO enzymes^41,47^. Like SUMO2/3, Smt3 has SUMOylation consensus sites in its unstructured N-terminal region that are utilized for chain formation (reviewed in ^48^), yet loss of the N-terminal lysines or the Smt3 N-terminus is tolerated without detectable growth defects^41^. The single *C. elegans* SUMO protein SMO-1 shares about 67% sequence identity with SUMO1 and 48% identity with SUMO2/3. Intriguingly, its N-terminus contains no lysine residues and can thus not contribute to canonical SUMO chain formation. As shown in Figures 1I an 1J, deleting the N-terminus in Smt3 and SMO-1 indeed enhanced interaction with SIM-containing model proteins, the SUMO E3 ligases *S.c.* Siz1^49^, and *C.e.* GEI-17^50^, respectively. These results suggest that the inhibitory effect of SUMO1’s N-terminus is indeed an evolutionarily conserved trait.

### SUMO1’s disordered N-terminus interferes with SIM-dependent functions

One explanation for increased interaction upon deletion of SUMO1’s N-terminus is that it usually occludes the SUMO1 SIM-binding groove, a feature apparently unique to SUMO1. If so, its removal should lead both to enhanced non-covalent interaction (Fig. 2A, C-F), and to enhanced SUMOylation (Fig. 2B, G-L).

**Figure 2.**
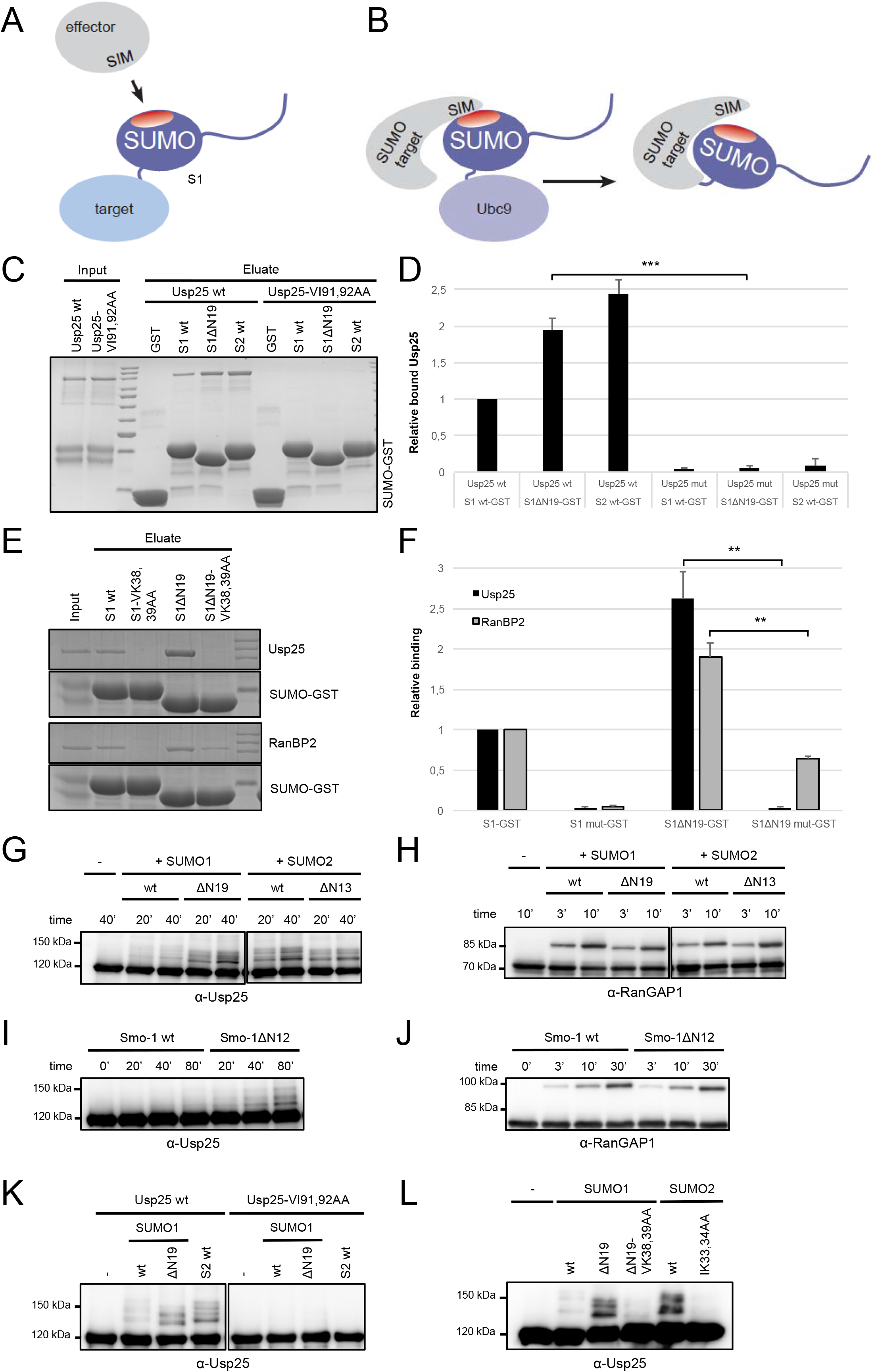
N-terminal deletions of SUMO1 augment binding and SUMOylation of Usp25 by enhancing SUMO-SIM interactions. (A) If not occupied by the N-terminus, the SIM-binding groove of SUMO1 can aid complex formation between SUMOylated proteins and their SIM-containing potential interaction partners. (B) SIMs in SUMOylation targets can promote SUMO-modification by stabilizing complex formation with SUMO-loaded Ubc9 termed “SIM-dependent SUMOylation”. (C) and (E) Pulldown of recombinant Usp25 wt, Usp25-V91A,I92A (SIM mutant), or RanBP2 with glutathione beads loaded with the indicated SUMO (S)-GST variants, analyzed by SDS-PAGE and Coomassie staining. (D) and (F) The amounts of Usp25 and RanBP2 in the eluates of SUMO pull-downs in (C) and (E) were quantified relative to SUMO-GST (n = 3). (G) - (L) *In vitro* SUMOylation reactions of recombinant Usp25 wt or -V91A,I92A, and RanGAP1 with the indicated human SUMO1, SUMO2 or *C. elegans* SMO-1 mutants, analyzed by SDS-PAGE and immunoblotting. Error bars represent one standard deviation. Asterisks indicate p-values: **: P ≤ 0.01; ***: P ≤ 0.001;

Indeed, a SIM-deficient USP25 (USP25 VI91,92AA^15^) lost binding not only to SUMO2 but also to SUMO1 ΔN19 (Fig. 2C + D), indicating that the interaction with truncated SUMO1 also depends on a bona fide SIM. Moreover, mutating two residues in SUMO1’s SIM-binding groove, Val38 and Lys39 (Fig. 2E + F) led to a significant loss of interaction with USP25 and with RanBP2.

We next turned to SUMOylation. As reported previously, USP25 was SUMOylated more efficiently with SUMO2 compared to SUMO1. However, SUMO1ΔN19 can SUMOylate USP25 as efficiently as SUMO2 (Fig. 2G). Consistent with binding assays (Fig. 1E-F), N-terminal truncation of SUMO2 did not further enhance SUMOylation efficiency. As a control, we used the SIM-independent SUMO target RanGAP1 (Fig. 2H), which was equally modified with all SUMO1 and SUMO2 variants. Similar results were obtained when we compared *C. elegans* wild-type SMO-1 and SMO-1ΔN12 for their ability to modify our model human substrates (Fig. 2I + J). In SUMOylation assays with USP25 SIM mutants (Fig. 2K) and with SUMO1ΔN19 variants that carried mutations in the SIM-binding groove (Fig. 2L), SUMOylation of USP25 was abolished, indicating that the increase in SUMOylation upon deletion of SUMO1’s N-terminus depends on a SIM-dependent interaction.

Taken together, our findings indicate that SUMO1’s intrinsically disordered N-terminus inhibits both SIM-dependent interactions and SIM-mediated SUMOylation. This offers an unanticipated explanation for the apparent SUMO2 preference of SIM-dependent SUMO binding partners such as USP25.

### NMR experiments indicate physical interaction between the N-terminus and the SIM-binding groove of SUMO1

To gain structural insights into the mechanism by which SUMO1’s N-terminus prevents SUMO - SIM interactions, we turned to nuclear magnetic resonance (NMR). To assess motions in the nanosecond/sub-nanosecond time regime, ^1^H-^15^N-SUMO1. Residues 21-93 exhibit uniform positive hNOEs, characteristic of stable, globular domains and thus consistent with the known folded structure of the core (Fig. 3A). The C-terminal residues of SUMO1 have hNOEs near or below 0, a hallmark of highly flexible regions and consistent with the high degree of flexibility commonly observed in Ubl proteins near their C-terminal Gly-Gly motif^39,51^. Residues 4-20 in the N-terminus of SUMO1 displayed lower hNOE values than those of the folded core consistent with their being more flexible. However, while residues 5-9 showed large negative hNOEs consistent with high flexibility, residues 10-20 have hNOE values close to 0. This is suggestive of a split in the SUMO1 N-terminus into a highly flexible region (residues 5-9) and a region of intermediate flexibility (residues 10-20) whose dynamics are slower but not as slow as the folded core. Residue 4 has a hNOE value close to 0 hinting at slower dynamics in the very N-terminal part of the protein. Unfortunately, residues 1-3 were not available for hNOE analysis, so no conclusions can be drawn about this sub-region of the N-terminus. The slower-than-expected dynamics of residues 10-20 indicate that this region spends time associated with the structured core. We speculate that the interactions could occlude the SIM-binding groove of SUMO1.

**Figure 3.**
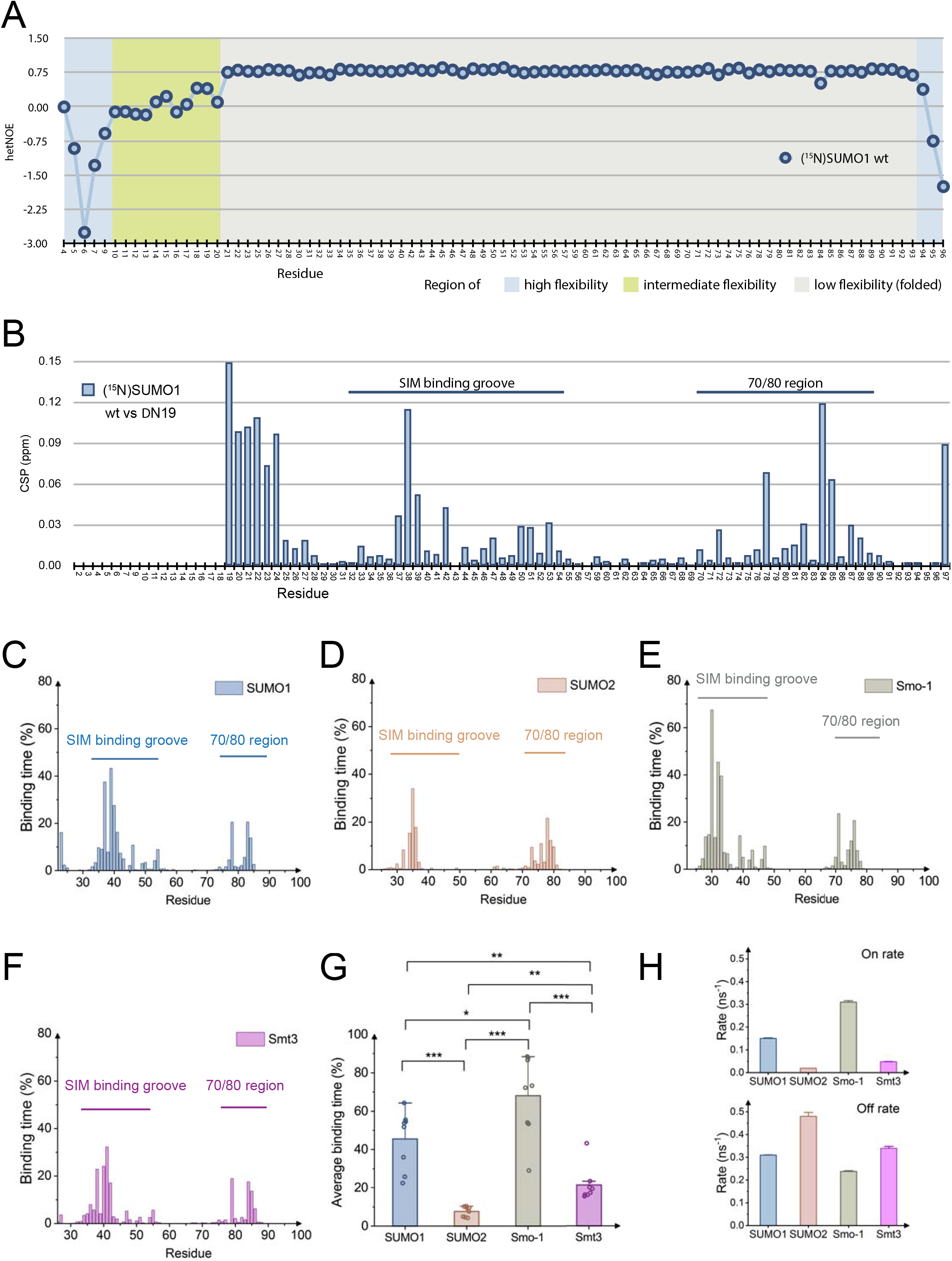
NMR experiment and Molecular Dynamics simulations reveal that the N- termini of SUMO1 and SMO-1 transiently occupy the SIM binding groove. (A) ^1^H-^15^N hNOE values for SUMO1. The x-axis shows SUMO1 residues in a non- contiguous fashion; residues not shown were either prolines (residues 8, 58 and 77) or could not be analyzed due to insufficient data quality (residues 1-3, 24, 29, 41 and 97). Colored backgrounds show areas with indicated dynamics. (B) HSQC-TROSY experiments comparing full-length SUMO1 wt and the ΔN19 variants. Histogram of CSPs of (^15^N)SUMO1 ΔN19 compared to full-length (^15^N)SUMO1; the resonance for Met19 could not be assigned confidently (shown as off-scale in histogram - marked with *); residues 1-18 are displayed only for clarity with a value of 0; Pro8, Pro58, Pro77 are not shown. The SIM binding groove and 70/80 region are marked (black bars). (C-F) Binding time, averaged over 8 1μs-trajectories, of the SUMO N-terminus to other residues for SUMO1, SUMO2, SMO-1 and yeast Smt3. (G) Average binding time of the SUMO N-terminus to the SIM binding groove for SUMO1, SUMO2, SMO-1 and yeast Smt3. Data points (n=8): average along single 1 μs trajectories; bars: average over all 8 trajectories, errors bars: standard errors of the mean of the 8 averages. Asterisks indicate p-values: *: P ≤ 0.05; **: P ≤ 0.01; ***: P ≤ 0.001. (H) Kinetic rate constants for binding/unbinding processes between the N-terminus of SUMO1, SUMO2, SMO-1 and yeast Smt3 and the SIM binding groove.

To investigate this further, we compared HSQC-TROSY spectra of full-length SUMO1 and the ΔN19 variant (Fig. 3B) to assess how the presence and absence of the N-terminus affects the structured core domain. Substantial chemical shift perturbations (CSPs) were observed for peaks in the core domain (CSPs above the 80^th^ percentile ranging from 0.032 to 0.119 ppm). If the N-terminus were flexible and structurally independent from the core domain, the spectra would be expected to be virtually identical. The CSPs appear as three clusters rather than randomly throughout the sequence: (1) residues 21-24 are directly adjacent to the flexible N-terminus and are thus expected to be affected when the latter is deleted, (2) residues 32-54 are in the SIM-binding groove, and (3) residues from the 70s to the 80s (dubbed here the “70/80 region”). Together with the hNOE results, the observations strongly suggest that the N-terminus interacts transiently with (at least) two specific regions of the core.

### MD simulations provide insight into SUMO1 N-terminus - core interactions

Given the timescale of the motions of the SUMO1 N-terminus as determined by hNOE, we turned to all-atom molecular dynamics (MD) simulations to gain further insights into the molecular mechanism by which SUMO1’s N-terminus prevents SUMO - SIM interactions. For extensive sampling of the conformational space accessible to the disordered N-terminal region, we initiated the MD simulation with different orientations of the N-terminus from the NMR structure for SUMO1 (see methods, Table S1). We used an IDP-adequate force field, and the simulations were validated by monitoring the Cα root-mean-square-deviation (RMSD) of the SUMO core against the initial structures (< 3 Å) (Fig. S2). In addition, chemical shifts back-calculated from MD simulations were in good accordance with the experimentally derived ones (Fig. S3).

In MD simulations, we observed the disordered N-terminus to interact reversibly with the SUMO core on the microsecond MD time scale, as measured by changes of the binding area between the N-terminus and the core (Fig. S4A + S4B and Table S1). The N-terminus of SUMO1 engaged with the core for about 65% of the simulation time resulting in an average minor shielding of the total core area of only 1.4% (Fig. S5A + S5B). Thus, the MD simulation, like the NMR experiments, points to a highly dynamic interaction of the SUMO1 N-terminus with the core. Moreover, MD simulations recapitulated findings by NMR by indicating that the SIM binding groove (residues His35, Lys37 and Lys39 primarily affected) and the 70/80 region are temporarily occupied by the SUMO1 N-terminus (Fig. 3C). Taken together, we conclude that MD simulations are a suitable predictive *in silico* tool to study the N-terminus/core

### MD simulations reveal significant differences of N-termini - core interactions across SUMO proteins

We next performed similar simulations for SUMO2, SMO-1 and Smt3. Again, we observed transient interactions of disordered N-termini of these paralogues with their respective cores on the microsecond MD time scale (Figs. 3D - G and S4C – S4E). However, comparison of all four SUMO proteins also revealed clear differences in average binding times and binding areas, with SMO-1 showing the longest binding time and largest binding area, followed by SUMO1, Smt3, and SUMO2 (Fig. 3G, Fig. S5A – S5D and Table S1). The most striking differences exist for the interaction with the SIM-binding groove, while interactions with the 70/80 region are rather comparable (Fig. 3C - F, compare Fig. S5B and 3G). These observations are consistent with a role of SUMO1 N-termini as intramolecular inhibitors of SUMO - SIM interactions, and show that this inhibitory effect is evolutionarily conserved in animals. The N-terminus of yeast Smt3, which is evolutionarily neither a SUMO1 nor SUMO2 protein (see ^41^ and above), shows an intermediate effect, while SUMO2’s N-terminus does not seem to inhibit SUMO - SIM interactions, all in congruence with our biochemical data (see Figs. 1 and 2).

### Interaction between SUMO’s disordered N-termini and the SIM binding groove is highly dynamic

Intrinsically disordered proteins exhibit particularly fast binding kinetics when binding does not involve conformational changes such as secondary structure formation^52,53^. SUMO’s N-termini form neither in the bound nor unbound states secondary structures in MD simulations consistent with PSIPRED secondary structure predictions^54^ indicating low local secondary structure propensities (Fig. S6A). Moreover, conformational adaption of the N-termini upon binding was very minor (Fig. S6B). We intrinsically disordered N-terminus to the SIM-binding groove. For this, we resorted to the MD simulations. We assumed a simple two-state model of bound and unbound states and determined the dwell times of all binding and unbinding events as measured by changes in binding area between the N-terminus of SUMO and the SIM binding groove (Fig. 3H and Fig. S5E). The MD-calculated off-rates koff for SUMO1, SUMO2, SMO-1, and yeast Smt3 range between 0.25 and 0.50 ns^-1^, thus are extremely high and hardly differ among the SUMOs. The on-rates vary by more than an order of magnitude, with the lowest on-rate for SUMO2 of 0.02 ns^-1^ and the highest on-rate for SMO-1 at 0.3 ns^-1^. While kon of SUMO2 and yeast Smt3 are significantly lower than koff resulting in an overall preference for the unbound state, the difference between on- and off rates is much less pronounced for SUMO1 (0.15 versus 0.31 ns^-1^) and for SMO-1 kon is higher than koff (0.31 versus 0.24 ns^-1^), signifying a preference for the bound state. In consequence, of the four N-termini, those of SUMO1 and SMO-1 possess the highest dwell time in the SIM-bound state in our simulations.

### Acidic residues in SUMO’s N-termini mediate the inhibitory effect on SIM-dependent interactions

SUMO1 residues that interacted most prominently in MD simulations with SUMO1’s N-terminus included His35, Lys37, Lys39, Lys46, and Arg54 (see Fig. 1B, 3C and 4A), suggestive of electrostatic interactions being vital for the interaction. All SUMO proteins have negatively-charged amino acids in their N-termini, albeit at different positions (see Fig. 1B, 4A, 4B and S6C). Most striking in SUMO1‘s N-terminus were two neighboring negatively-charged residues, Glu11 and Asp12. Indeed, mutating these residues to Lys individually or jointly to disrupt the N-terminus-SIM-binding groove interaction strongly increased the interaction of SUMO1 with USP25 and with TDP2 (Fig. 4C + 4D) as well as SIM-dependent SUMOylation (Fig. 4G + H).

**Figure 4.**
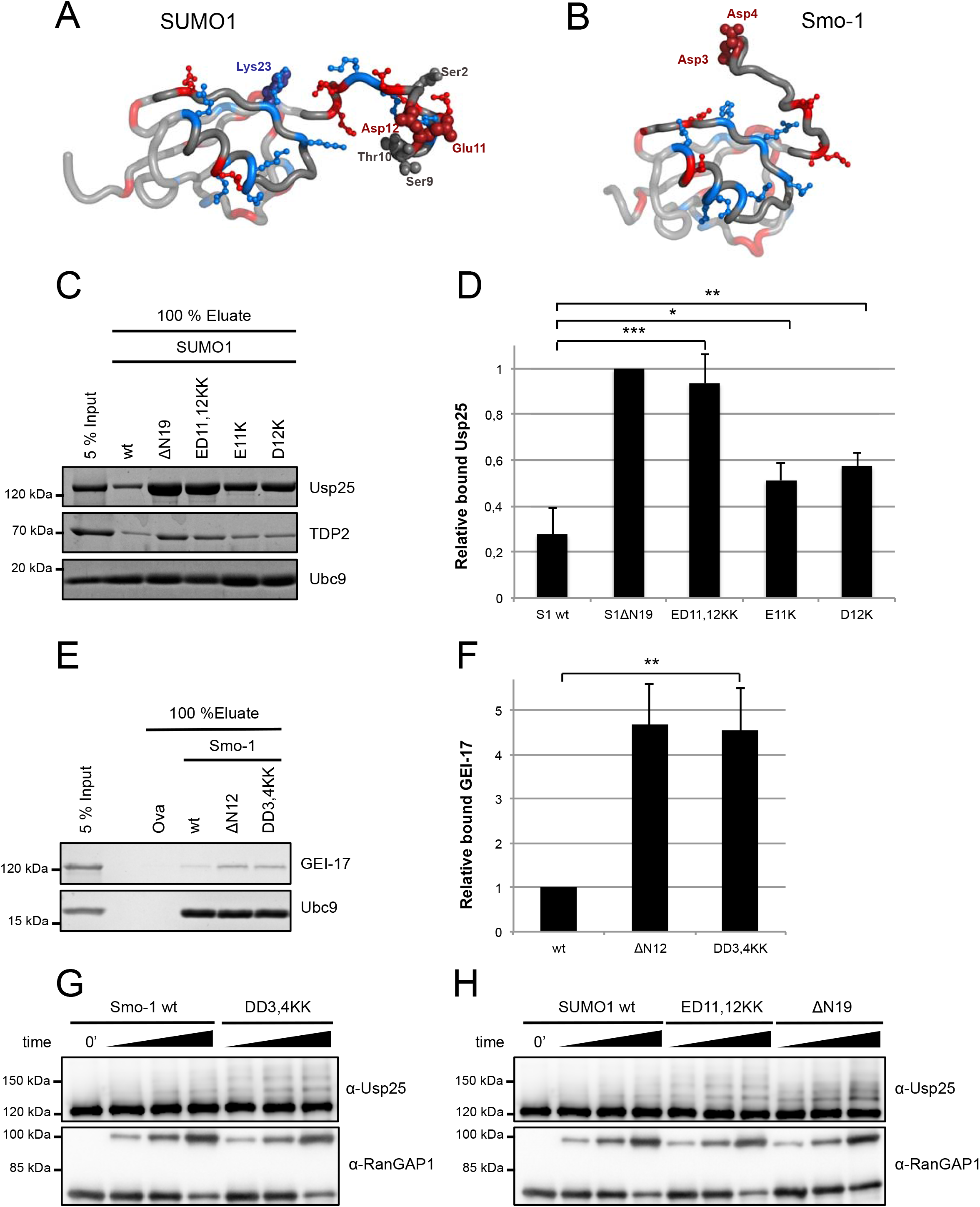
Mutating negatively charged residues in the N-terminus of SUMO1 abolishes the inhibitory effect of the N-terminus. The SIM binding groove, encompassing β2-strand and α-1 helix, and N-terminus were shown as non-transparent. The positively/negatively charged residues were colored in blue/red, respectively. The charged residues in N-terminus and around the SIM binding groove were shown as ball and stick. The residues related with post-translational modifications or mutations were highlighted in the figure with their names. (A) human SUMO1; (B) *C. elegans* SMO-1; (C) Pulldown of recombinant Usp25, Ubc9 or TDP2 with the indicated SUMO1 beads analyzed by SDS-PAGE and Coomassie staining. (D) The amounts of Usp25 in the eluates of SUMO pull-downs in (C) were quantified relative to Ubc9 (n = 6). (E) Pulldown of recombinant *C. elegans* GEI-17 or hUbc9 with Ovalbumin (Ova) or indicated SMO-1 beads analyzed by SDS-PAGE and Coomassie staining. (F) The amounts of GEI-17 were quantified relative to hUbc9 (n = 3). (G) and (H) *In vitro* SUMOylation of recombinant Usp25 and RanGAP1 with human SUMO1 wt or the indicated mutants, analyzed by SDS-PAGE and immunoblotting. Error bars represent one standard deviation. Asterisks indicate p-values: *: P ≤ 0.05; **: P ≤ 0.01;

Next, we tested whether our findings with human SUMO1 also hold true for *C. elegans* interaction with the SIM-containing SUMO E3 ligase GEI-17 (Fig. 4E + F). Taken together, our findings suggest that the intrinsically disordered N-termini of SUMO1 proteins inhibit SIM-dependent interactions and SUMOylation due to electrostatic interactions of N-terminal acidic residues with basic residues in SUMO’s core.

### The SUMO1 N-terminus shields the SIM-binding groove via ionic interactions

To substantiate interpretations from the biochemical assays described above, we compared the HSQC-TROSY NMR spectrum of a ^15^N-labeled SUMO1 ED11,12KK variant with that of wild-type SUMO1. Strikingly, there are large CSPs in the entire N-terminus and in the region of the core directly adjacent to the N-terminus (Fig. 5A). The N-terminal CSPs are more widespread than would be predicted for amino acid changes in an IDR, suggesting that the mutant N-terminus experiences a different chemical environment to the wild-type. There are small, yet notable CSPs in the SIM-binding groove. This finding confirms that the ED11,12KK mutations disrupt the interaction between the N-terminus and the SIM-binding groove and stem. Consistent with this notion, MD simulations of SUMO1 ED11,12KK reveal that the mutations significantly decrease interaction with the SIM-binding groove (Fig. 5 B - D). Similar behavior was observed comparing wt *C. elegans* SMO-1 with SMO-1 DD3,4KK (Fig. 4B, 5E - G). The slight increase in interactions with the 70/80 region seen in the MD simulations of both mutants, however, can not be supported by the CSPs (Fig. 5A), and is potentially caused by the known overstabilization of salt bridges in canonical protein force fields^55^. Together, our findings indicate that the SUMO1 N-terminus acts as a cis inhibitor of SUMO - SIM interactions due to its ability to mask the SIM binding groove via dynamic ionic interactions.

**Figure 5.**
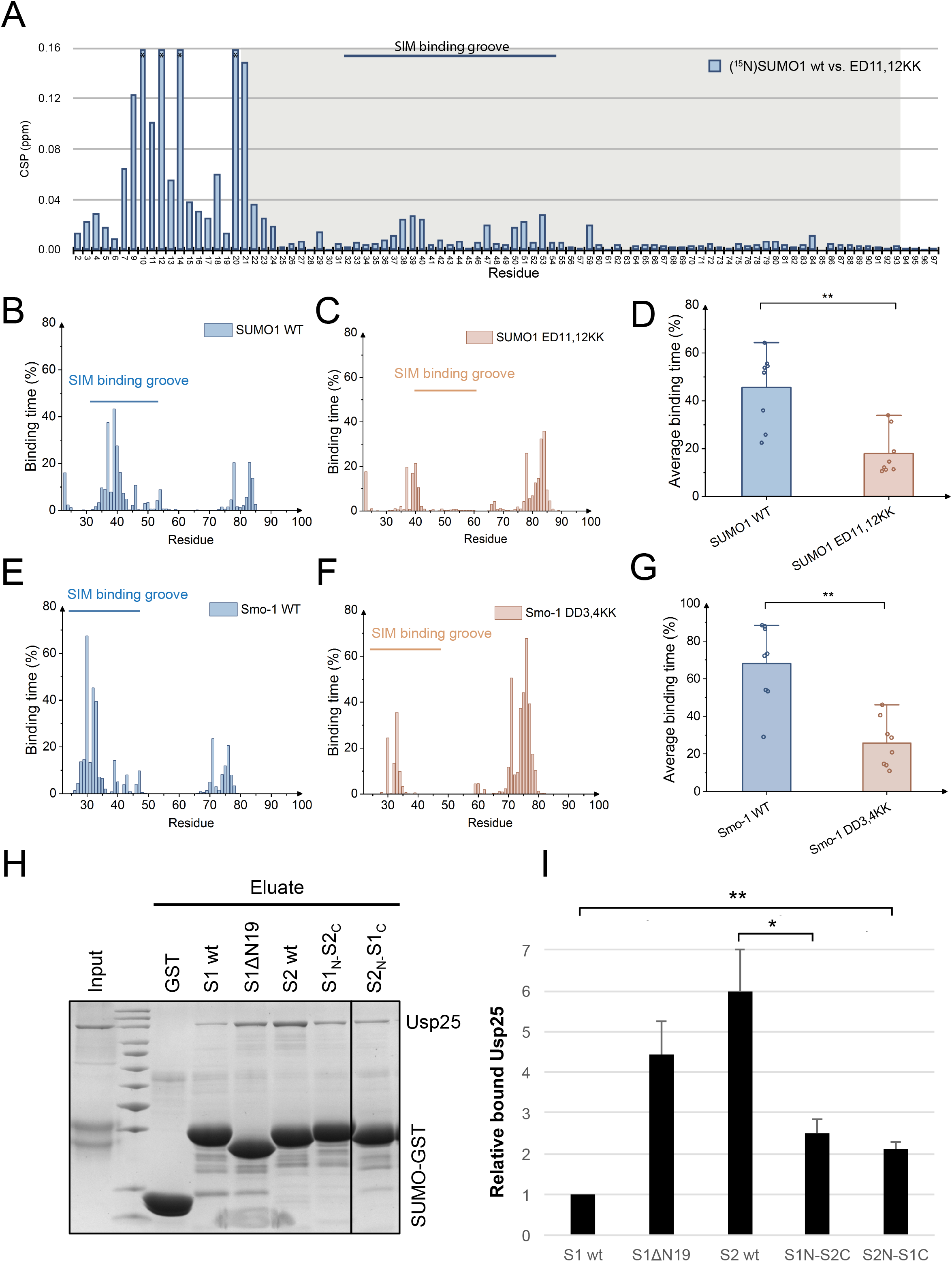
Negatively charged residues in the N-terminus of SUMO1 mediate the inhibitory effect on SIM-dependent interactions (A) HSQC-TROSY experiments comparing full-length SUMO1 wt and the ED11,12KK mutant. Histogram of CSPs of (^15^N)SUMO1-ED11,12KK compared to full-length (^15^N)SUMO1; the resonances for Thr10, Lys12, Gly14 and Glu20 of the mutant could not be assigned confidently (shown as off-scale in histogram - marked with *); Pro8, Pro58 and Pro77 are not shown. The SIM binding groove (black bars) as well as the folded core of the protein (grey background) are marked. (B-C) Binding time and (D) Average binding time of the SUMO N-terminus to the SIM binding groove for wt (identical to Fig. 3C) and ED11,12KK variant of SUMO1. (E-F) Binding time and (G) Average binding time of the N-terminus of wt (identical to Fig. 3E) and DD3,4KK of SMO-1 to residues of the SUMO core. Data points: average along single 1 μs trajectories; bars: average over 8 trajectories, errors bars: standard errors of the mean of the 8 averages. (H) Pulldown of recombinant Usp25 with GST-tagged SUMO1/SUMO2 chimeras, analyzed by SDS-PAGE and Coomassie staining. (I) The amounts of Usp25 in the eluates of SUMO pull-downs in (H) were quantified relative to the SUMO-GST proteins (n = 3). Asterisks indicate p-values: *: P ≤ 0.05; **: P ≤ 0.01

### The inhibitory function of SUMO1’s N-terminus can be partially transplanted to SUMO2

The inhibitory effect of SUMO1‘s N-terminus seems largely driven by ionic interactions with residues flanking the SIM-binding groove (Fig. 4A + B). SIM-binding grooves are flanked by basic residues in all SUMO paralogues, raising the question of whether its effect could be transplanted onto SUMO2. We created several SUMO1/SUMO2 chimeras and tested them for interaction with USP25 in pulldown assays (Fig. 5H - I). Intriguingly, a SUMO2 chimera with SUMO1’s N-terminus (S1N-S2C) showed reduced interaction. Moreover, a SUMO1 core that carries the SUMO2 N-terminus (S2N-S1C) bound USP25 more efficiently than wt SUMO1, albeit not as efficiently as SUMO2. In conclusion, SUMO N-termini are transplantable elements that contribute to the differences of SUMO proteins in SIM - dependent interactions.

### Phosphorylation may regulate the inhibitory effect of SUMO’s intrinsically disordered N-termini

Based on the ionic nature of the interaction, it is conceivable that posttranslational modifications that add or remove charges may influence inhibition. Numerous SUMO residues have been identified by proteomics as acetylated, phosphorylated, or methylated (summarized on phosphosite.org), some of which have been validated (e.g., pS2 in SUMO1^56^, or AcK32 in SUMO2^57^).

To explore these ideas further, we turned again to MD simulations. We tested an array of modifications for SUMO1 (Fig. 6A - C and S7A – E), of which phosphorylation of Ser9 alone or in combination with Thr10 significantly increases the residence time of the SUMO1 N-terminus on the SIM-binding groove predicting that phosphorylation at these sites may further inhibit SIM-dependent interactions. Whether these predictions hold true and translate into a functional outcome needs to be tested experimentally in the future using post-translationally modified forms of SUMO.

**Figure 6.**
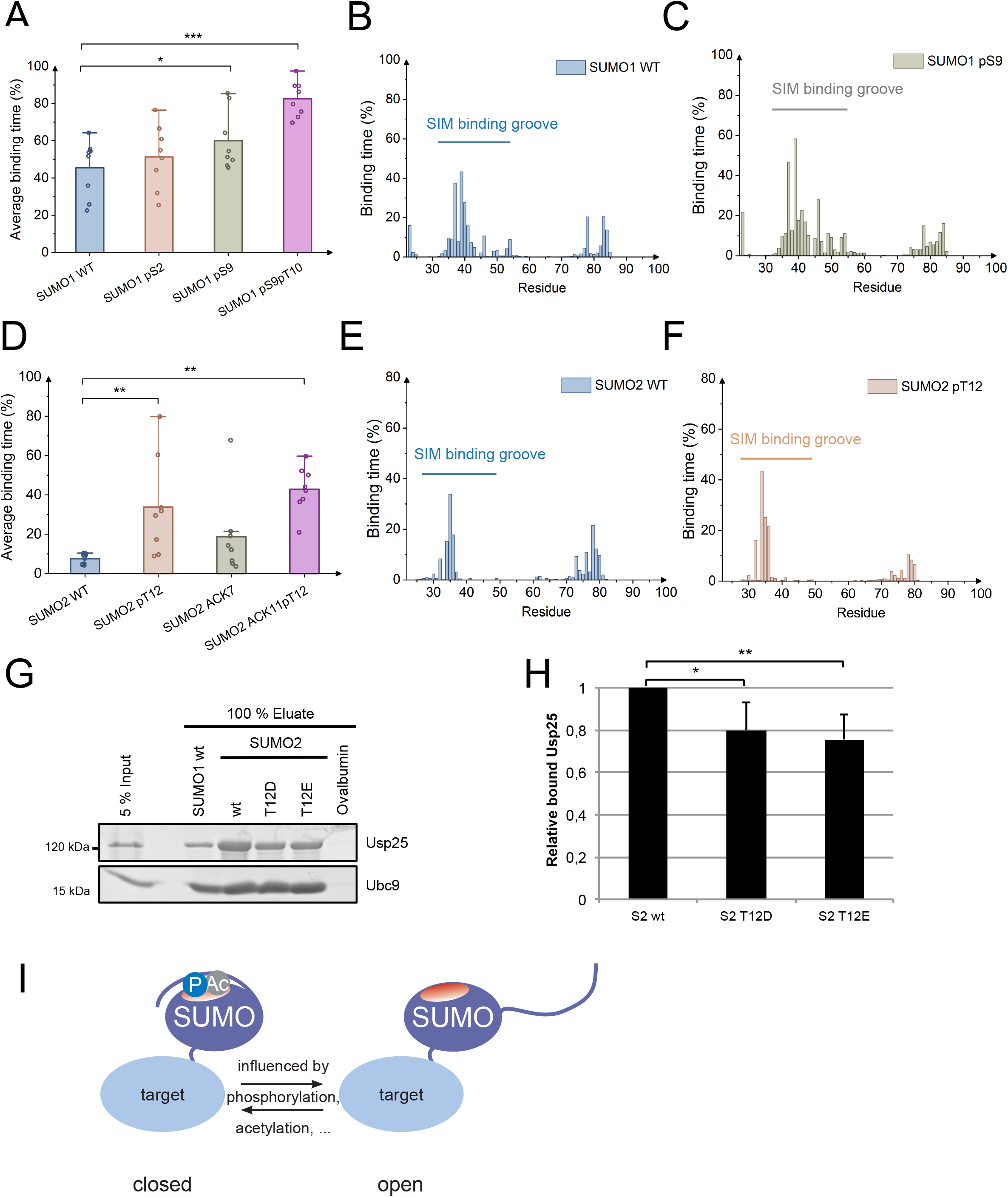
Post-translation modifications of SUMO’s intrinsically disordered N- termini may regulate SIM-dependent interactions. (A) Average binding time of the SUMO N-terminus to the SIM binding groove for SUMO1 wt (identical to Fig. 3C) and pS2, pS9, and pS9pT10 SUMO1 variants. (B) and (C) Binding time of the N-terminus of SUMO1 wt (identical to Fig. 3C) and SUMO1-pS9 to individual residues of the SUMO core. (D) Average binding time of the SUMO N- terminus to the SIM binding groove for SUMO2 wt and the pT12, AcK7 and AcK11pT12 SUMO2 variants. (E) and (F) Binding time of the N-terminus of SUMO2 wt (identical to Fig. 3D) and SUMO2-pT12 to individual residues of the SUMO core. Data points: average along single 1 μs trajectories; bars: average over 8 trajectories, errors bars: standard errors of the mean of the 8 averages. (G) Pulldown of recombinant Usp25 or Ubc9 with the indicated SUMO1 beads, analyzed by SDS-PAGE and Coomassie staining. (H) The amounts of Usp25 in the eluates of SUMO pull-downs in (G) were quantified relative to Ubc9 (n = 4). Error bars represent one standard deviation. Asterisks indicate p-values: *: P ≤ 0.05; **: P ≤ 0.01; ***: P ≤ 0.001. (I) Model of the intrinsically disordered region of SUMO serving as a regulatable cis - inhibitor for SIM - dependent interactions through post-translation modification.

For SUMO2 (Fig. 6D - F and S7F – S7J), MD simulations indicate a significant increase in the overall residence time on the SIM-binding groove upon phosphorylation of Thr12. Could the increased interaction of the N-terminus with the SIM-binding groove predicted by MD simulation lead to a reduction of SIM-dependent interactions? In the absence of quantitatively phosphorylated SUMO2, we turned to phospho-mimetic variants. Indeed, mutating Thr12 in SUMO2‘s N-terminus either to Asp or Glu significantly decreased the interaction of SUMO2 with USP25 (Fig. 6G + H). Together, these findings allow speculation that the N-terminus of SUMO2 may act as an inhibitor of SUMO - SIM interactions upon posttranslational modification(s).

In conclusion, we reveal a function for SUMOs‘ intrinsically disordered N-termini as intramolecular regulators of SUMO-dependent interactions. N-termini of human and *C. elegans* SUMO1 proteins are clearly inhibitory and other SUMO N-termini may acquire such a function upon posttranslational modification of the N-terminus (Figure 6I).

### Deletion of the flexible N-terminus in *C. elegans* SMO-1 reduces germ cell survival

Finally, we wanted to address the biological significance of our findings. The N-terminus of yeast Smt3 only shows an intermediate effect on SIM binding (our findings above), serves as an acceptor for SUMO chains, and is dispensable for normal growth^40,41^. We thus turned to *C. elegans* that expresses the single SUMO protein SMO-1. Its N-terminus is not only a strong inhibitor of SIM interactions (our findings described above), it is also devoid of lysines and can thus not contribute to canonical SUMO chain formation.

We created an in-frame deletion allele by CRISPR/Cas9 genome editing in *C. elegans smo-1* (*smo-1(zh156)*, referred to as SMO-1ΔN12), removing 11 amino acids at the N-terminus but leaving the start methionine. SMO-1ΔN12 animals were viable and displayed no obvious anatomical defects or embryonic lethality. We could also not detect significant differences in SUMO levels and overall SUMOylation in total animal extracts (Fig. S8). However, SMO-1ΔN12 mutant hermaphrodites had a smaller brood size (i.e. the average number of progeny per animal) than wild-type controls suggesting a defect in germline development or oocyte fertilization (Fig. 7B). The *C. elegans* sumoylation pathway is important for germ cell development and specifically, for chromosome congression in germ cells at the pachytene stage of meiosis I^50,58,59^. Whole-mount DAPI staining revealed that the gonads of SMO-1ΔN12 mutants contained a smaller pachytene region with fewer germ cell nuclei but a similar density of nuclei (i.e. number of nuclei per area) as wild-type controls (Fig. 7A, C - E). This phenotype could be due to decreased germ cell proliferation or to increased germ cell apoptosis. Under physiological conditions, around half of the germ cells at the pachytene stage undergo apoptosis, and the rate of apoptosis can be further increased by introducing environmental stress such as DNA damage^60^. We therefore monitored germ cell apoptosis in SMO-1ΔN12 mutants. First, we examined the CED-1::GFP reporter, which outlines apoptotic corpses that are engulfed by the somatic sheath cells^61^, and observed an increased number of CED-1::GFP corpses in adult SMO- 1ΔN12 animals 24 and 48 hours post L4 stage despite the overall reduced number of pachytene cells (Fig. 7F + 7G). As a second assay, we performed live-staining with SYTO12, a dye that accumulates in germ cells undergoing apoptosis^61^. Also SYTO12 staining revealed an elevated number of apoptotic germ cells in adult SMO-1ΔN12 mutants 29 hours post L4 stage (Fig. 7H). To determine whether the increased levels of apoptosis in the SMO-1ΔN12 mutant are due to elevated DNA damage, we used the *cep-1(gk138)* p53 loss-of-function allele, which causes resistance to DNA-damage-induced germ cell apoptosis. SMO-1ΔN12; *cep-1(gk138)* double mutants showed levels of apoptosis comparable to WT or *cep-1(gk138)* single mutants (Fig. 7G). Thus, the increased apoptosis levels in SMO-1ΔN12 mutants depend on the DNA damage response.

**Figure 7:**
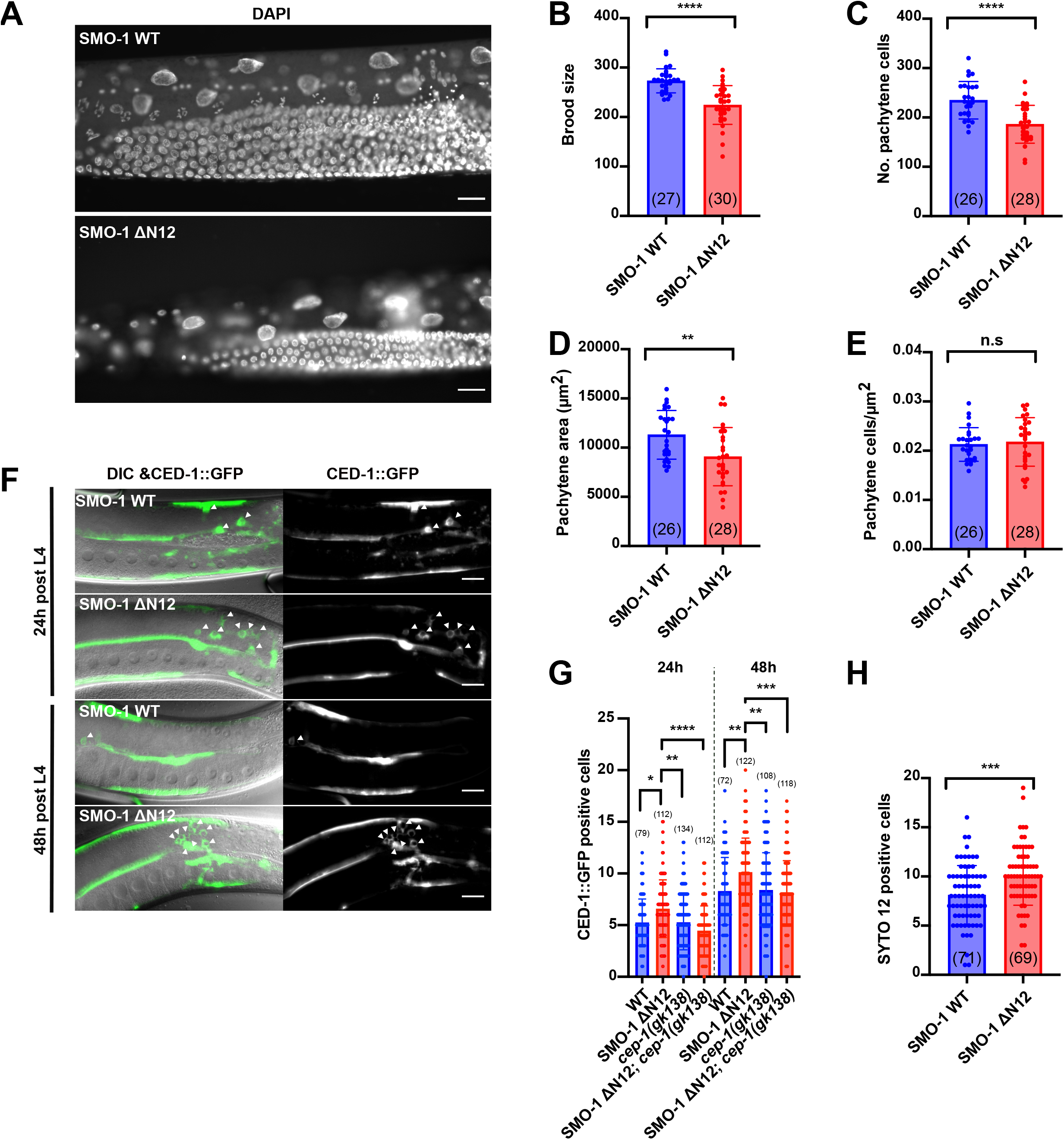
Deletion of the flexible N-terminus in C. elegans SMO-1 reduces germ cell survival. (A) Whole-mount DAPI staining of the wild-type and of SMO-1ΔN12 (smo-1(zh156)) animals 24 hours post L4 stage. (B) Brood size, (C) number of germ cell nuclei at the pachytene stage per gonad arm, and (D) area of the pachytene region in wild-type and SMO-1ΔN12 animals. (E) Average number of pachytene stage germ cell nuclei per area. (F) Apoptotic corpses outlined by CED-1::GFP staining in wild-type and SMO- 1ΔN12 mutants 24 and 48 hours post L4. The engulfed corpses are indicated with white arrowheads. The left panels show the DIC images overlayed with CED-1::GFP signal (average intensity projections of the outer gonad layers) and the right panels the CED- 1::GFP signal alone. (G) Number of CED-1::GFP-positive apoptotic corpses per gonad arm in wild-type and SMO-1ΔN12 animals with or without *cep-1(gk148)* background 24 and 48 hours post L4 stage. (H) Number of SYTO12 positive germ cells per gonad arm in wild-type and SMO-1ΔN12 animals 29 hours post L4 stage. Bars indicate average values, dots individual values and error bars the standard deviations. The numbers in brackets indicate the numbers of animals scored. If the values were normally distributed (in B - E, H), statistical significance was calculated with an unpaired parametric t-test, or else (in G) a Kruskal-Wallis test was used. Asterisks indicate the p-values as * p≤0.05; ** p≤ 0.01; *** p≤ 0.001; **** p≤0.001, n.s. for not significant. The scale bars in (A & F) represent 20µm.

Together, these data indicate that the flexible N-terminus of *C. elegans* SMO-1 is required for normal germline development and in particular for the survival of germ cells entering the pachytene stage of meiosis.

## DISCUSSION

### SUMO N-termini are Intrinsically <u>D</u>isordered <u>R</u>egions

Intrinsically disordered proteins and protein regions (IDP/IDR) have been largely overlooked until the 1990s even though they constitute 40% of the proteome of eukaryotes^62^ and fulfill many important biological functions including signal transduction^63,64^. Both their mechanism of action and biological functions vary greatly. For example, IDP/IDRs may function as (auto)inhibitors of enzymes or act as assembly platforms for multi-subunit complexes (reviewed in ^65^).

Unstructured N-terminal regions of 13 - 23 amino acids are a hallmark of all SUMO proteins (Fig. 1B, Table S1) and distinguishes them from Ubiquitin and most other Ubl relatives. These feature the expected amino acid composition of IDRs^66^, namely a relatively low content of hydrophobic amino acids along with a higher ratio of polar or charged amino acids (Figure 4A + 4B, and S6). The disordered and flexible nature of SUMO N-termini function has remained rather enigmatic. Some SUMO N-termini engage in chain formation (e.g., ^67,68^). Additionally, SUMO N-termini have been suggested to act as entropic bristles that may prevent aggregation, but SUMO aggregation has only been observed upon extended incubation at very high temperature^69^. Here, we discovered a rapid nanosecond equilibrium between "open" SUMO conformations available for interactions with SIMs and "closed" ones that are cis-inhibited by the N-terminal region. The N-terminal region remains fully disordered in the bound state during our MD simulations and is thus a classic example of intrinsic disorder irrespective of the binding state.

### SUMO N-termini form fuzzy complexes with SUMOs’ SIM binding grooves

How specificity is achieved by an IDR at the molecular level is an important question. Upon binding to their biological partners, IDRs often undergo a conformational transition to a folded state, also referred to as coupled binding and folding^70^. A well-known example is the cell cycle inhibitor p27, which folds around cyclin / Cdk complexes^71^. Nevertheless, increasing evidence demonstrates that at least some IDRs remain unstructured upon interaction with partners, which is referred to as formation of a "fuzzy complex"^72^. Interactions in such a complex are mediated by transient hydrophobic and electrostatic interactions in the fuzzy region, which can provide non-specific contacts or specific interactions with multiple motifs^53^. Examples include the fuzzy complex of transcriptional activator Gcn4 and its partner Gal11/Med15^73^, ultrafast interaction between FG-repeat containing nucleoporins and nuclear transport receptors^74^, or the fuzzy intramolecular binding between the IDR and folded domain of Src kinase^75^ or galectin^76^.The interaction mode of SUMOs’ N-termini with the SUMO core is in line with the fuzzy complex model. First, we observed in MD simulations a very fast (ns) rate of the binding / unbinding process, and secondly, the bound state of the SUMO N-terminus and the SUMO core is transient and dynamic. Although the N-terminus hovers specifically over the SIM binding groove, it does not assume a specific stable configuration but rather forms a "ligand cloud" (Fig. S6B)^77^. Despite this fuzzy nature, the N-terminus can compete efficiently with SIM-containing proteins for the SIM binding groove, as supported by both experiments and simulations.

### SUMO N-termini are cis-inhibitors of SIM-dependent interactions

The N-termini of recombinant SUMO1, SMO-1 and Smt3 act as inhibitors of SIM dependent interactions (this study,^42^), and potentially also of interactions with other SUMO surfaces, namely the 70/80 region. A recent study by Lussier-Price et al. also reported on a paralogue-specific auto-inhibition of SIM binding by SUMO1’s N- terminus^42^. Similar to our study in the outcome but with a different experimental approach, they show strongly reduced KDs for the phosphorylated SIMs of PML and Daxx upon deletion of SUMO1’s but not SUMO2’s N-terminus and shielding of a hydrophobic area on SUMO1 by the N-terminus that can be competed off by a phosphorylated SIM peptide of PML.

Inhibition via a module within the same protein (cis-inhibition) plays an important role in regulation^78^. IDRs are frequently involved in cis-inhibition, possibly as a way to economize genome/protein resources^65^. A consequential benefit of this mechanism is the ease of regulation via posttranslational modifications^79^. This is likely the case for SUMOs‘ N-termini because the interaction of SUMOs‘ IDRs with the SIM-binding grooves is largely mediated by electrostatics, which may be strengthened or weakened by a host of posttranslational modifications that are emerging for SUMO proteins both in the IDR (e.g., ^56,68^ or flanking the SIM binding groove^34,57^). Here we provide the first evidence for this idea from MD simulations and pulldown assays (Fig. 6 and Fig. S7). An important implication from our analyses is that the SUMO2 N-terminus may become a cis-inhibitor upon phosphorylation of Thr12.

### SUMO N-termini add control to SIM-dependent interactions

SIM-dependent interactions contribute to all aspects of the SUMO pathway yet how specificity can be ensured with a single SUMO binding module is still unclear. Increasing evidence suggests regulation of SUMO - SIM interactions at multiple levels. For example, acetylation of lysine residues surrounding the SIM binding groove can alter the affinity for specific binding partners (see, e.g., ^57,80^), while phosphorylation of residues in a given SIM may increase the affinity for SUMO and change SUMO paralogue preferences (e.g., ^33,35^). Our findings add yet another layer of regulation: whether SUMO on a given target is available for interaction with a downstream effector can be controlled via SUMO‘s intrinsically disordered N-terminus (model in Fig. 6I). Our analysis of *C. elegans* SMO-1 points to a specific function of the SUMO N-terminus in effects in the germline and soma, affecting a variety of cell fates, deletion of SMO-1’s N-terminus reveals increased p53-dependend germ cell apoptosis, suggesting a specific role of the SMO-1 N-terminus in controlling DNA damage-induced apotosis. These findings are consistent with previous reports showing that the SUMO system in *C. elegans* plays an essential role in sensing environmental stress and maintaining genome stabiblity^81^. Taken together, our findings offer an intriguing explanation for the presence and strict evolutionary conservation of SUMOs’ intrinsically disordered N-termini: they seem to equip SUMO with an environment-sensing and regulatory capacity of SIM-dependent processes including SIM-dependent modification and downstream effector binding.

### Limitations of the study

Here we assigned an important function to SUMO1 and Smo-1 N-termini as inhibitors of SUMO-dependent interactions, and provide an explanation for paralogue preferences of SUMO targets such as USP25. An exciting question that we discuss is whether the inhibitory potential of different SUMO - N-termini can be tuned by posttranslational modifications of the N-termini or the SUMO core (model in Fig. 6I). While our MD simulations and *in vitro* studies with selected mutants point in this direction, we have not been able to generate quantitatively acetylated and/or phosphorylated SUMO variants to test this hypothesis. An important question for future studies will be the identification of pathways and protein targets for which this specialized function of SUMO’s N-terminus is most relevant. Although Smt3’s N-terminus can be deleted without detectable effect on growth^41^, at least in standard laboratory conditions, we are not aware of any eukaryotic organism with a SUMO protein that lacks the flexible N-terminus. In light of our findings *in C. elegans*, were deletion of Smo’s N-terminus increases p53-dependent cell death, we speculate that stress response pathways that provide populations with evolutionary advantages are the right places to look for.

## STAR+METHODS

Detailed methods are provided in the online version of this paper and include the following:

## KEY RESOURCES TABLE

Lead contact and materials availability statement METHOD DETAILS

SUMO nomenclature Expression constructs

Protein expression and purification In vitro binding assays

In vitro SUMOylation assays Quantification and Statistical Analysis

Structure preparation and Molecular Dynamics (MD) simulation system setup Energy minimization, equilibration and molecular dynamics simulations Analyses of molecular dynamics simulation

## STAR+METHODS

### Key Resources Table

Contact for Reagent and Resource Sharing

Further information and requests for reagents may be directed to, and will be fulfilled by the corresponding author Frauke Melchior.

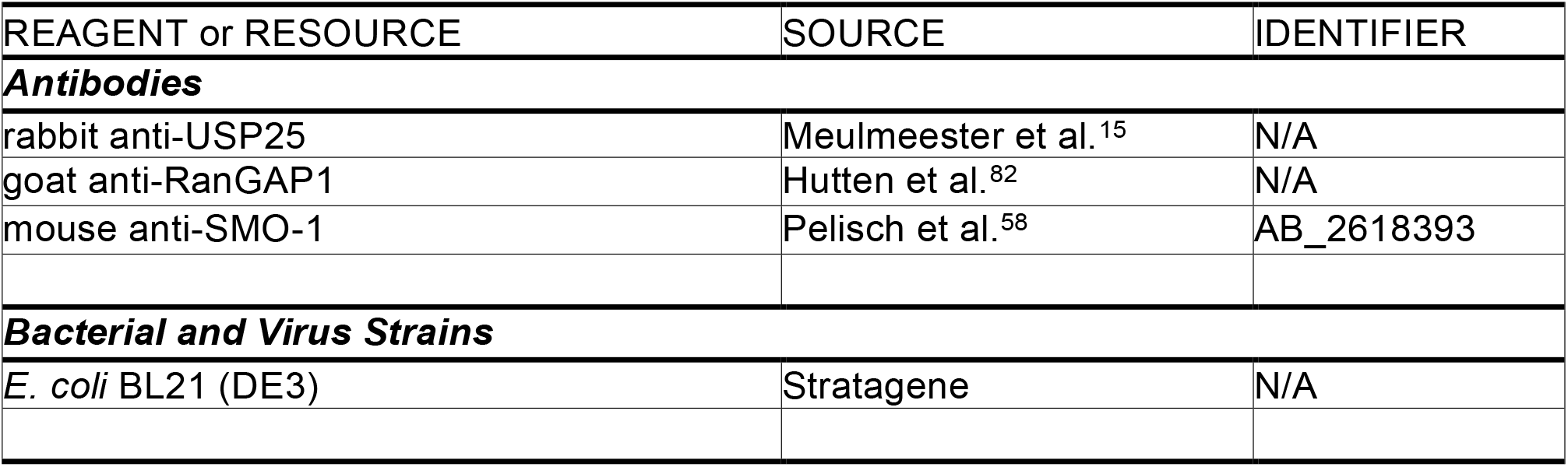
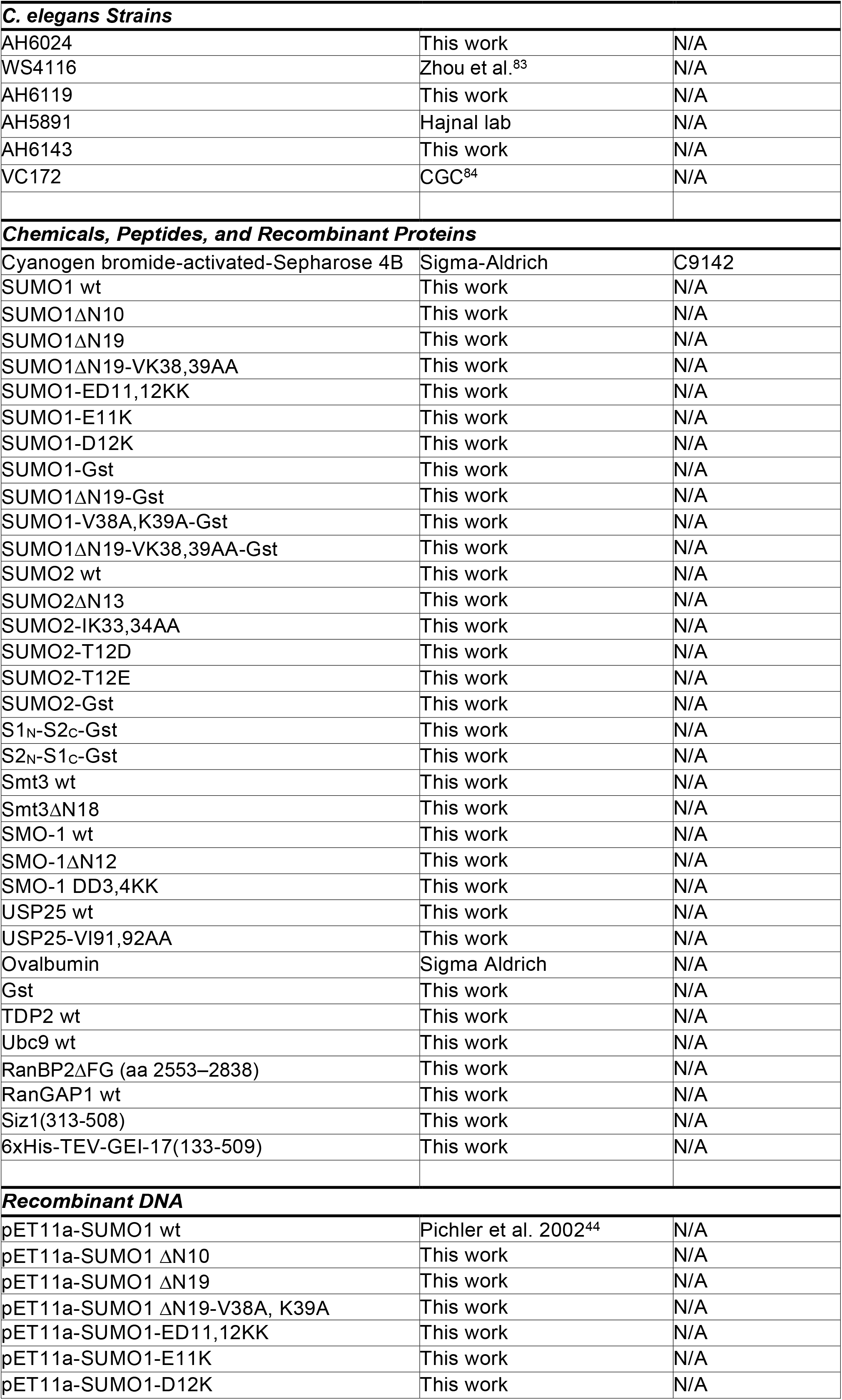
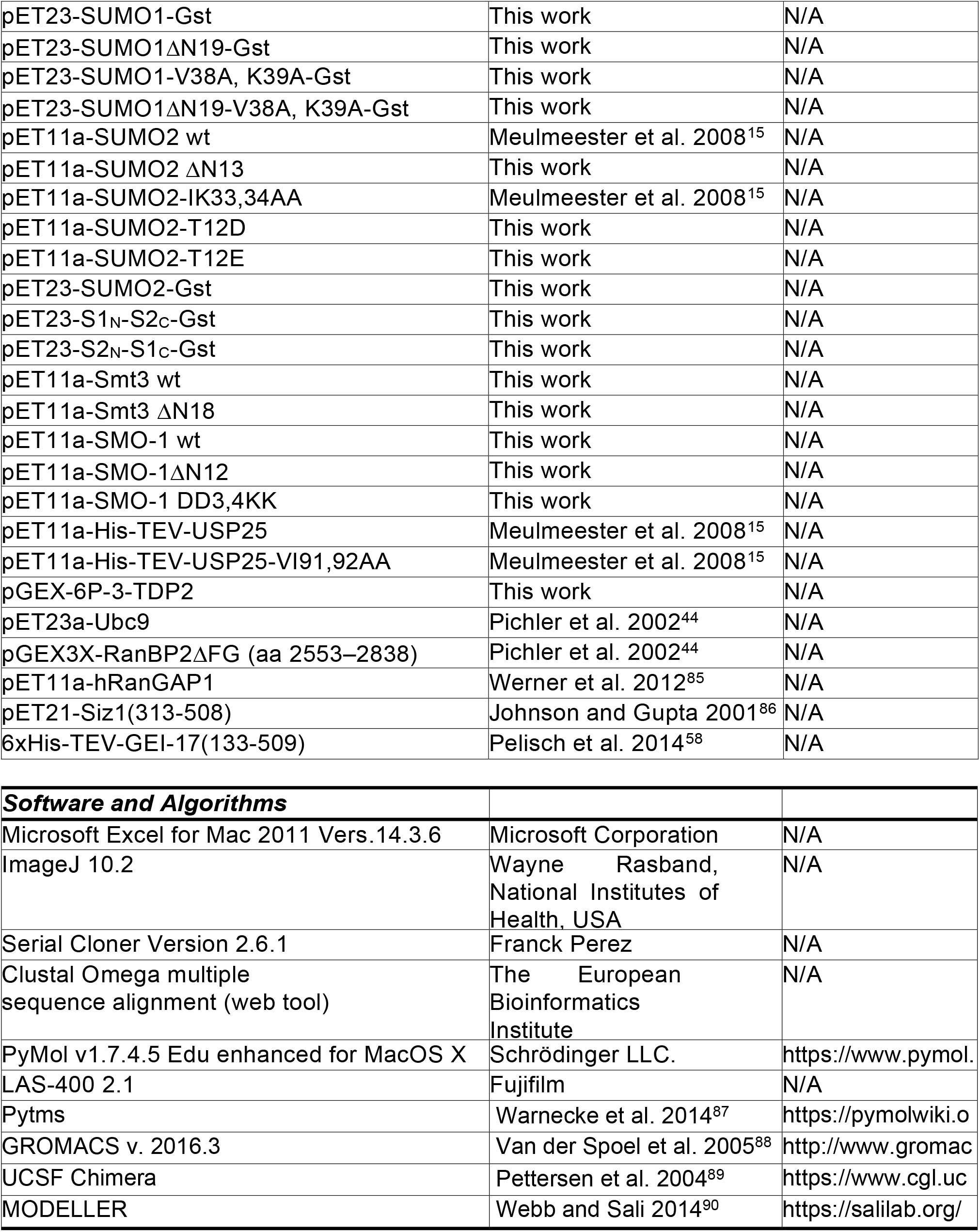

## Method Details

### SUMO nomenclature

The nomenclature for mammalian SUMO2 and SUMO3 is used inconsistently. Like many colleagues in the SUMO field, we follow the nomenclature as introduced by ^91^.

Their assignment was consistent with the original description of mammalian SUMO genes (reviewed in ^3^). According to this, mature SUMO2 (Smt3A) is 92 amino acids long, mature SUMO3 (Smt3B) consists of 93 amino acids.

### Expression constructs

Bacterial expression constructs for untagged SUMO1(ΔC4) and SUMO2(ΔC11), USP25, USP25-V91A, I92A^15^, GST-RanBP2 ΔFG, Ubc9, SAE1 (Aos1), SAE2 (Uba2)^44^ and RanGAP1^85^ were previously described. All constructs but one were based on human cDNA sequences. The Ubc9 construct was derived from mouse cDNA, but the encoded mouse protein is identical to human Ubc9. The coding sequence of mature full-length *S. cerevisiae* Smt3 (Smt3ΔC3) and *C. elegans* SMO-1 (SMO-1ΔC1) were PCR-amplified and cloned into the NdeI and BamHI sites of pET11a. The coding sequence of mature full-length human SUMO1 and SUMO2 was PCR-amplified and Gibson-cloned into the SacI and NotI sites of pET23a; GST was fused to the C terminus of SUMO by PCR-amplification of GST and cloning into the NotI and XhoI sites. SUMO1ΔN19 and SUMO2ΔN13 were PCR-amplified and cloned into the NdeI and BamHI sites of pET11a. Other N-terminal deletions and amino acid mutations of SUMO-, Smt3-, and SMO-1 plasmids were performed using the QuikChange site-directed mutagenesis method (Agilent) or by Gibson DNA assembly. For the SUMO1/SUMO2 hybrid constructs, the SUMO1 and SUMO2 N- and C-termini (SUMO1 N: aa 1-20, C: aa 21-97; SUMO2 N: aa 1-15, C: aa 16-92) were PCR-amplified from the C-terminal GST fusion constructs including roughly half of the vector backbone each and fused in the desired combinations by Gibson DNA assembly. The coding sequence of human TDP2 was PCR-amplified from the pENTR221-TDP2 plasmid (genomics and proteomics core facility, DKFZ) and cloned into the BamHI and XmaI sites of the pGEX-6P-3 vector. The expression plasmid for the catalytic fragment of Siz1(313-508)-pET21 was a kind gift by Dr. Erica S. Johnson^86^. 6xHis-TEV-GEI- 17(133-509) was a gift from Ronald Hay^58^ (Addgene plasmid #87056; http://n2t.net/addgene:87056 ; RRID:Addgene_87056). The sequences of all expression constructs used in this work were verified by Sanger sequencing.

### Protein expression and purification SUMO proteins

Untagged SUMO1-, SUMO2-, Smt3- and SMO-1 variants were essentially purified according to an established protocol^92^. *E. coli* BL21 (DE3) cells were transformed with the respective pET11a-SUMO plasmid and a single colony was grown over night in LB medium supplemented with ampicillin. The cells were centrifuged for 5 min at 3900 g and resuspended in fresh LB medium supplemented with ampicillin or MOPS medium supplemented with (^15^N)ammonium chloride and ampicillin (for NMR). When the main culture reached an OD600 of 0.6 - 0.8, expression of SUMO variants was induced with 1 mM IPTG and expressed for 3 - 4 h at 37°C or overnight at 16°C . Cells were harvested by centrifugation for 10 min at 7200 g, resuspended in 50 mM Tris–HCl pH 8.0, 50 mM NaCl, 1 mM DTT, supplemented with 1 μg/ml of each aprotinin, leupeptin, pepstatin A and flash frozen in N2(l). After thawing, the cell suspension was lysed using an EmulsiFlex-C5 (Avestin) in the presence of DNase I and cleared by ultra-centrifugation for at least 45 min at 100,000 g and 4 °C. The supernatant was incubated with 10 ml DEAE- or Q-Sepharose for at least 1 h at 4 °C, the sepharose material was sedimented and the supernatant was concentrated using a centrifugal filter unit with 5 kDa MWCO. The protein was then purified by gel filtration over a Superdex200 column equilibrated in Transport Buffer (20 mM HEPES-KOH pH 7.3, 110 mM KAcO, 2 mM Mg(AcO)2, 1 mM EGTA, 1 mM DTT) supplemented with 1 μg/ml of each aprotinin, leupeptin, pepstatin A (TB+++). In some but not all SUMO preparations, an additional gel filtration run in 50 mM Tris pH 7.5, 1M LiCl, 150 mM NaCl, 1 mM DTT, 1 μg/ml of each aprotinin, leupeptin, pepstatin A was included before the gel filtration in TB+++. SUMO containing fractions were pooled, concentrated and applied to a Superdex75 gel filtration column equilibrated in TB+++. SUMO containing fractions were pooled, concentrated, flash-frozen in N2(l) and stored at -80 °C.

All SUMO-GST proteins were expressed from the respective pET23a-SUMO-GST plasmids and lyzed in 50 mM Tris-HCl pH 8.0, 100 mM NaCl, 1 mM EDTA, 1 mM EGTA, 1 mM CaCl2, 1 mM MgCl2, 1 mM DTT, 1 μg/ml of each aprotinin, leupeptin, pepstatin A, Pefa in the presence of DNAase I as above, and the protein in the supernatant after ultracentrifugation was purified by pull-down with glutathione agarose equilibrated in GST lysis buffer. Bound protein was washed with GST wash buffer (50 mM Tris-HCl pH 8.0, 300 mM NaCl, 1 mM EDTA, 1 mM DTT, 1 μg/ml of each aprotinin, leupeptin, pepstatin A) and eluted with GST wash buffer supplemented with 20 mM glutathione. SUMO-GST containing fractions were pooled, concentrated and applied to a Superdex75 gel filtration column equilibrated in TB+++. SUMO-GST containing fractions were pooled, diluted 1:2 with 50 mM Tris pH 7.5, bound to a MonoQ column equilibrated in 50 mM Tris pH 7.5. Bound protein was eluted with a gradient of 50 mM –1 M NaCl in 50 mM Tris pH 7.5. SUMO-GST containing fractions were pooled, concentrated, flash-frozen in N2(l) and stored at -80 °C.

### GST-tagged TDP2 and RanBP2ΔFG

For expression of N-terminally GST-tagged TDP2 and RanBP2ΔFG, *E. coli* BL21 (DE3) cells were transformed with the pGEX-6P-3-TDP2 and the pGEX-3X-RanBP2ΔFG plasmid, respectively, and a single colony was grown over night in LB medium supplemented with ampicillin. The cells were centrifuged for 5 min at 3900 g and resuspended in fresh LB medium supplemented with ampicillin. When the main culture reached an OD600 of 0.6 - 0.8, it was shifted to 16 °C, expression was induced with 0.5 mM IPTG and expressed for 21 h at 16 °C. Cells were harvested by centrifugation for 10 min at 7200 g, resuspended in GST lysis buffer and flash frozen in N2(l). After thawing, the cell suspension was lysed using an EmulsiFlex-C5 and cleared by ultra- centrifugation for at least 45 min at 100,000 g and 4 °C. The GST-tagged protein in the supernatant was then purified by pull-down with glutathione-sepharose equilibrated in GST lysis buffer. For this, the supernatant was incubated with 10 ml glutathione sepharose / l main culture for at least 1 h at 4 °C. The bead material was then collected in a column and washed with 50 CV GST lysis buffer. Elution was performed with GST elution buffer. Protein-containing fractions were pooled, concentrated and applied to a Superdex200 gel filtration column equilibrated in TB+++. Protein containing fractions were pooled, concentrated, flash frozen in N2 (l) and stored at - 80 °C.

### USP25

His-TEV-USP25 was transformed into *E. coli* BL21 (DE3) and a single colony was grown over night in LB medium supplemented with ampicillin. The cells were centrifuged for 5 min at 3900 g, resuspended in fresh LB medium supplemented with ampicillin and grown at 37 °C. When the main culture reached an OD600 of 0.6 - 0.8, it was shifted to 30 °C, USP25 expression was induced with 0.5 mM IPTG and expressed for 4 h. Cells were harvested by centrifugation for 10 min at 7200 g, resuspended in 50 mM Na-phosphate pH 8.0, 300 mM NaCl, 10 mM imidazole, 1 mM DTT, 1 μg/ml of each aprotinin, leupeptin, pepstatin A and flash-frozen in N2 (l). The cells were lysed using an EmulsiFlex-C5. The supernatant after 100,000 g ultracentrifugation was incubated with Ni-NTA Sepharose for 2 h at 4 °C and eluted with 50 mM Na-phosphate pH 8.0, 300 mM NaCl, 200 mM imidazole, 1 mM DTT, 1 μg/ml of each aprotinin, leupeptin, pepstatin A. Protein-containing fractions were pooled, concentrated and applied to a Superdex200 gel filtration column equilibrated in TB+++. USP25 in the eluate was pooled, concentrated, flash-frozen in N2(l) and stored at -80 °C.

### RanGAP1

Untagged RanGAP1 was expressed and purified according to an established protocol^93^. *E. coli* BL21 (DE3) cells were transformed with pET11a-RanGAP1 and a single colony was grown over night in LB medium supplemented with ampicillin. The cells were centrifuged for 10 min at 5000 g and resuspended in fresh LB medium supplemented with ampicillin, 1 mM MgCl2 and 0.1 % glucose. When the main culture reached an OD600 of 0.8 - 0.9, RanGAP1 was induced with 0.5 mM IPTG and expressed for 3 h at 37 °C. Cells were harvested by centrifugation for 15 min at 5000 g, resuspended in 50 mM Tris–HCl pH 8.0, 100 mM NaCl, 1 mM EDTA, 1 mM DTT, supplemented with 1 μg/ml of each aprotinin, leupeptin, pepstatin A and flash frozen in N2(l). After thawing, the suspension was supplemented with 1 mg/ml lysozyme, incubated for 1 h on ice and lysed by sonication (from this point all buffers supplemented 1 μg/ml of each aprotinin, leupeptin, pepstatin A). The pellet after centrifugation for 30 min at 20,000 g was washed by resuspension using a glas douncer and centrifugation successively in 50 mM Tris–HCl pH 8.0, 1% Triton X-100, and in 50 mM Tris–HCl pH 7.4, 2 M urea. The final pellet was solubilized in 50 mM Tris–HCl pH 7.4, 8 M urea. The supernatant was dialyzed against 50 mM Tris–HCl pH 7.4, 100 mM NaCl with two buffer exchanges, centrifuged at 100,000 g for 1 h, applied to a Q–Sepharose (GE Healthcare) column equilibrated with 50 mM Tris–HCl pH 7.4, 150 mM NaCl, washed with 50 mM Tris–HCl pH 7.4, 300 mM NaCl and eluted with 50 mM Tris–HCl pH 7.4, 1 M NaCl. Protein fractions were then applied to a HiPrep- 16/60 Sephacryl S-300 HR (GE Healthcare) column equilibrated in TB+++ and RanGAP1- containing fractions were concentrated, pooled, flash-frozen in N2 (l) and stored at -80 °C.

### SUMO E1 activating enzyme (SAE1-SAE2 heterodimer)

Expression and purification of the SAE1-SAE2 heterodimer followed an established protocol^44^. *E. coli* BL21 (DE3) cells were co- transformed with pET28a-Aos1 (His-SAE1) and pET11d-Uba2 (SAE2) and grown for 18 h at 37 °C in 500 ml LB medium supplemented with kanamycin and ampicillin. After addition of 1.5 l of LB medium supplemented with kanamycin and ampicillin, protein expression was induced with 1 mM IPTG and performed for 6 h at 25 °C. Cells were harvested by centrifugation for 15 min at 5000 g, resuspended in E1 lysis buffer (50 mM Na-phosphate pH 8.0, 300 mM NaCl, 10 mM imidazole) and flash-frozen in N2(l).

After thawing, the respective cell suspensions were supplemented with 1 μg/ml of each aprotinin, leupeptin, pepstatin A and 1 mM DTT. Cells were lysed using an EmulsiFlex- C5. The supernatant after ultracentrifugation at 100,000 g for 1 h was incubated with with E1 wash buffer (E1 lysis buffer with 20 mM imidazole) and eluted with E1 elution buffer (E1 lysis buffer with 250 mM imidazole). Protein-containing fractions were pooled, concentrated and applied to a Superdex200 gel filtration column equilibrated in TB+++. Eluate fractions were checked for the presence of both proteins, His-SAE1 and SAE2, by SDS-PAGE. The respective fractions were pooled and applied to a MonoQ 5/50 GL column (GE Healthcare) equilibrated in 50 mM Tris–HCl, pH 7.5, 50 mM NaCl, 2 mM DTT, 1 μg/mL of each aprotinin, leupeptin, pepstatin. The E1-complex was eluted by applying a linear gradient from 50 to 500 mM NaCl in 20 column volumes. Collected fractions were analyzed by SDS-PAGE and complex containing fractions were pooled, dialyzed against TB+++ and flash-frozen in N2(l).

### SUMO E2 conjugating enzyme Ubc9

Expression and purification of Ubc9 followed an established protocol^92^. *E. coli* BL21 (DE3) cells were transformed with pET23a-Ubc9 and a single colony was grown over night at 37 °C in LB medium supplemented with ampicillin, 1 mM MgCl2 and 0.1 % glucose. The cells were centrifuged for 10 min at 5000 g and resuspended in fresh LB medium supplemented with ampicillin, 1 mM MgCl2 and 0.1 % glucose. When the main culture reached an OD600 of 0.6, Ubc9 expression was induced with 1 mM IPTG and expressed for 3 h at 37 °C. Cells were harvested by centrifugation for 15 min at 5000 g, resuspended in Ubc9 lysis buffer (50 mM Na-phosphate buffer, pH 6.5 supplemented with 1 mM DTT, 1 μg/ml of each aprotinin, leupeptin, pepstatin A) and flash-frozen in N2(l). After thawing, the lysate was cleared by 100,000 g ultracentrifugation at 4 °C and applied to an SP-Sepharose column equilibrated in Ubc9 lysis buffer. The column washed extensively using Ubc9 lysis buffer and Ubc9 eluted with Ubc9 lysis buffer supplemented with 300 mM NaCl. Protein-containing fractions were applied to a Superdex75 gel filtration column equilibrated in TB+++. Ubc9 in the eluate was pooled, concentrated, flash-frozen in N2(l) and stored at -80 °C.

### SIM-containing fragment of the SUMO E3 ligase S. c. Siz1 (aa 313-508)

Siz1(313-508)-pET21 was transformed into *E. coli* BL21 (DE3) and a single colony was grown over night in LB medium supplemented with ampicillin. The cells were centrifuged for 5 min at 3900 g, resuspended in fresh LB medium supplemented with ampicillin and grown at 37 °C. When the main culture reached an OD600 of 0.6 - 0.8, Siz1 expression was induced with 1 mM IPTG and expressed for 4 h. Cells were harvested by centrifugation for 10 min at 7200 g, resuspended in 50 mM Na-phosphate pH 8.0, 300 mM NaCl, 10 mM imidazole, 1 mM DTT, 1 μg/ml of each aprotinin, leupeptin, pepstatin A and flash-frozen in N2 (l). After thawing, the cells were lysed using an EmulsiFlex-C5. The supernatant after 100,000 g ultracentrifugation was incubated with Ni-NTA Sepharose for 2 h at 4 °C and eluted with 50 mM Na-phosphate pH 8.0, 300 mM NaCl, 200 mM imidazole, 1 mM DTT, 1 μg/ml of each aprotinin, leupeptin, pepstatin A. Protein-containing fractions were pooled, concentrated and applied to a Superdex75 gel filtration column equilibrated in TB+++. Siz1 in the eluate was pooled, concentrated, flash-frozen in N2 (l) and stored at -80 °C.

### SIM-containing fragment of the SUMO E3 ligase C. e. GEI-17 (aa 133-509)

Rosetta2 (DE3) cells transformed with 6xHis-TEV-GEI-17 were grown in LB medium supplemented with ampicillin, 1 mM MgCl2, 0.1% glucose at 37°C. Expression was induced with 0.1 mM IPTG for 16 h at 20°C. The cells were harvested, resuspended in lysis buffer (50 mM Tris-HCl pH 7.5, 500 mM NaCl, 10 mM imidazole, 0.1% Triton- X100, 1 mM DTT, 1 μg/ml of each aprotinin, leupeptin, pepstatin A, Pefa bloc) and flash-frozen in N2 (l). After thawing, the cells were lysed using an EmulsiFlex-C5. After 100,000 g ultracentrifugation the supernatant was passed over a Ni-NTA sepharose column at 4 °C, the column was first washed with lysis buffer, then with 50 mM Tris-HCl pH7.5, 500 mM NaCl, 30 mM Imidazole, 1 mM DTT, 1 μg/ml of each aprotinin, leupeptin, pepstatin A. Bound protein was eluted with 50 mM Tris-HCl pH 7.5, 150 mM NaCl, 200 mM imidazole, 1 mM DTT, 1 μg/ml of each aprotinin, leupeptin, pepstatin A. Protein-containing fractions were pooled, concentrated and applied to a Superdex200 gel filtration column equilibrated in TB+++. GEI-17-containing elution fractions were pooled, concentrated, flash-frozen in N2 (l) and stored at -80 °C.

### *In vitro* binding assays

For the preparation of SUMO sepharose, SUMO1, SUMO2, Smt3 or SMO-1 concentrations were determined thoroughly by absorption at 280 nm using the protein-specific absorption coefficients, and by Bradford assay. The concentration measurements were verified by SDS-PAGE followed by Coomassie staining. Importantly, we only compared SUMO proteins prepared in the same way, either including or excluding the additional LiCl purification step in one experiment. Equal amounts of the untagged SUMO proteins were coupled to Cyanogen bromide-activated-Sepharose in Carbonate buffer (0.2 M pH 8.9) at a concentration of 1 mg protein per ml beads^15^. To block remaining coupling sites, beads were subsequently incubated with 100 mM Tris-HCl pH 8.0. Beads were washed several times with PBS Buffer (TB: 20 mM HEPES/KOH pH7.3, 110 mM potassium acetate, 2 mM magnesium acetate, 0.5 mM EGTA, 1 mM DTT and protease inhibitors) supplemented with 1 mM sodium azide.

Equal amounts of the SUMO-GST variants were bound to Glutathione sepharose in PBS supplemented with 0.05% Tween20 at a concentration of 1 mg/ml for 1 h at 4°C. Beads were washed several times with PBS supplemented with 0.05% Tween 20.

The coupling efficiency of all SUMO beads was always controlled by comparing the protein content of the coupling reaction before and after coupling.

For binding assays, 10 μg of recombinant binding partner per 300 μl reaction was incubated with 10 - 20 μl (USP25, TDP2, RanBP2, Siz1, GEI-17) or 5 - 7.5 μl (Ubc9) SUMO beads in TB supplemented with 0.05% Tween 20, and 0.2 mg/ml ovalbumin for at least 1 h at 4 °C. Beads were washed three times with TB or TB supplemented with 0.05% Tween 20 and eluted with 2x SDS sample buffer. The samples were analyzed by SDS-PAGE followed by Coomassie staining. The gels were scanned using the gel documentation system LAS-4000 (GE Healthcare) and the respective protein containing bands quantified with ImageJ.

### *In vitro* SUMOylation assays

*In vitro* SUMOylation assays with purified recombinant proteins were essentially performed as described previously^92^. Reactions containing USP25 were prepared with 100 nM USP25, 50 nM SAE1/SAE2 (E1 enzyme), 500 nM Ubc9 (E2 enzyme), 5 μM SUMO or SMO-1 variants, and 1 or 5 mM ATP in TB supplemented with 0.05% Tween 20 and 0.2 mg/ml ovalbumin and incubated at 30 °C for up to 80 minutes. Reactions containing RanGAP1 were prepared with 500 nM RanGAP1, 10 nM SAE1/SAE2, 10 nM Ubc9, 5 μM SUMO or SMO-1 variants, and 1 or 5 mM ATP and incubated at 30 °C for up to 30 minutes. Reactions were stopped by adding SDS sample buffer and incubation at 95 °C for 5 min. Samples were analyzed by SDS-PAGE followed by immunoblotting.

### Quantification and Statistical Analysis of biochemical experiments

The amount of SUMO binding partners in the eluates of SUMO or Smt3 binding reactions were quantified relative to the respective Ubc9 or SUMO-GST signal, if applicable, using ImageJ^94^. Data were analyzed using a paired Student’s t-test in Excel. Error bars represent one standard deviation. The number of independent experiments (n) is given in the respective figure legend. Asterisks indicate p-values: *: P ≤ 0.05; **: P ≤ 0.01; ***: P ≤ 0.001; n.s.: not significant.

### NMR spectroscopy

NMR spectra were recorded on a 500 MHz Bruker Avance II (University of Washington) at 295 K in 25 mM sodium phosphate pH 7.0, 150 mM NaCl, 10% D2O. ^1^H/^15^N-HSQC-TROSY and hNOE experiments were collected with 370 µM labeled protein (700 µM for SUMO1 ΔN19). Datasets were processed using NMRPipe/NMRDraw^95^, and visualized with NMRView^96^. Chemical shift perturbations observed by 2D HSQC-TROSY NMR were quantified in parts per million with the equation Δδ(^1^H/^15^N) = {[δ(^1^H) - δ(^1^H)0]^2^ + 0.04[δ(^15^N) - δ(^15^N)0]^2^}^1/2^.

### Structure preparation and Molecular Dynamics (MD) simulation system setup

The initial structures for wild type SUMO1 were selected from NMR structures with the PDB codes 1A5R^39^ and 2N1V^97^, and for mature wild type SUMO2 (92 aa) from an NMR structure with the PDB code 2RPQ^38^ without any preference for the distance between the N-terminal tail and the main body of SUMO.

The initial structure for yeast Smt3 was truncated from the X-ray structure of yeast Smt3 in complex with ULP1 (PDB code: 1EUV^98^). The missing N-terminal and C-terminal parts were modelled by MODELLER^90^. The initial structure for C. elegans SMO-1 were randomly selected from NMR structures with the PDB code 5XQM^99^. The mutants of SUMO were prepared using UCSF Chimera^89^ based on the wild type structures. Structures for a) SUMO1 that was acetylated on Lys23 (ACK23), b) SUMO1 that was phosphorylated on Ser2/Ser9/Ser9Thr10 (SUMO1 pS2/pS9/pS9pT10), and c) SUMO2 that was phosphorylated/acetylated on Lys7 (SUMO2 ACK7), Thr12 (SUMO2 pT12) and Lys11Thr12 (SUMO2 ACK11pT12) were prepared by using PyTMs^87^, a PyMOL^100^ plugin. The forcefield parameters of phosphorylated residues and acetylated Lys were reported previously^101,102^.

### Energy minimization, equilibration and molecular dynamics simulations

Molecular Dynamics simulations were performed using the GROMACS 2016.3 package^88^. The system was described by the Amber99sb*-ILDN force field^103^. Each SUMO structure was immersed in an explicit TIP4PD^104^ truncated dodecahedron water box, with at least 2.5 nm between periodic replicas of the protein (about 940 - 1540 nm^3^). The system was neutralized by adding ions, and extra NaCl was added to represent a solution with an ionic strength of 0.15 M to mimic physiological conditions. The systems contained 125 to 200k atoms and were minimized using the steepest descent minimization approach. After the minimization, the system was equilibrated in the NVT ensemble for 200 ps with all heavy atom position-restrained with a force thermostat with a coupling constant of 0.1 ps. Further equilibration was conducted in the NPT ensemble with all heavy atoms position-restrained for 200 ps and Cα atoms position-restrained for 1 ns, where the pressure was maintained at 1 atmosphere using a Parrinello-Rahamn barostat with the coupling constant set to 2.0 ps.

All equilibrations were performed with a time step of 1 fs. For the production run, the thermostat and barostat settings were the same as for the NPT run. To enable 2 fs time steps, all bonds were constrained to equilibration length using the LINCS algorithm^105^. A real-space cutoff of 10 Å was used for the electrostatic and Lennard-Jones forces. Snapshots from each trajectory were saved to disk every 50 ps. The simulations performed in this work are listed in Table S1.

### Analyses of molecular dynamics simulation

The change of solvent-accessible surface area (SASA) was used to quantify the binding and unbinding process between the N-terminus and the SUMO SIM binding groove or the complete SUMO core (Fig. S3 and S9). In case of the SIM binding groove, the ‘unbound’ state includes all conformations with the N-termini not interacting with the SUMO core and interacting with the core outside of the SIM-binding motif. Kinetic rate constants were obtained by fitting the integrated dwell binding/unbinding times. Six residues in the N-terminus adjacent to the SUMO core were not included for calculating the contact area between the N-terminus and the SUMO core or the SIM binding groove, respectively, to leave enough space (about 2 nm) for the SASA calculation probe (radius of the probe is 0.14 nm) (Fig. S3A). A threshold (0.3 nm^2^) was used in binding event determinations with the SUMO core or the SIM binding groove. The percentage of binding time of the N-terminus with the SIM binding groove or the SUMO core was calculated as the ratio of MD snapshots with binding areas (ΔSASA) larger than 0.3 nm^2^ along MD simulations. The analyses were using GROMACS utilities, and the first 100 ns of all production trajectories were regarded as equilibration and not used in final analyses. The MD back-calculated chemical shifts were calculated by using SHIFTX2^106^. The structures in the figures were prepared using PyMol^100^.

### General methods and maintenance of *C. elegans*

The Caenorhabditis elegans strains were maintained at 20°C on the standard NGM (Nematode Growth Medium) plates seeded with E. coli bacterial strain OP50 as a food source^107^. The derivate of Bristol strain N2 served as a wild-type reference. To generate double mutants, standard crossing methods were used. The list of used strains in this study is provided in the Key Resources Table and including the genotype in Table S2.

### CRISPR/Cas9 genome editing

Genome editing was performed according to the co-CRISPR strategy described by^108^. An oligonucleotide corresponding to a target sequence near the smo-1 translational start site (sgRNA #1: 5‘ GCCGATGATGCAGCTCAAGC 3‘) was cloned into the plasmid pMW46 (derivate of pDD162 from Addgene). The deletion of the eleven amino acids ADDAAQAGDNA at the SMO-1 N-terminus was achieved using the oligonucleotide pAF64 as repair template: CTC TAC CTC TCT CCT TCT ATC TCT TTT TCT CTT TTC AAA TCT AAT TTC GTT TCA GAG ACT CCC GCT ATA AAC GAT GGA ATA CAT CAA GAT CAA GGT CGT TGG ACA GGT AAT TTG ACT GGA AAT

TCG CCG CGA ATT TGT TAA TAA TTC CC. The following constructs with indicated final concentration were microinjected into young adult N2 hermaphrodites: dpy-10 sgRNA PJA58 – 25 ng/µl, dpy-10 donor template oligonucleotide AF-ZF-827 – 0.5 nM, smo-1 sgRNA #1 pAF25 – 75 ng/µl, smo-1 donor template oligonucleotide pAF64 – 0.5 nM and the transformation marker myo-2::mCherry pCFJ90 – 2.5 ng/µl. Two to three days post injection, 100 F1 recombinants showing a Rol phenotype were transferred to separate NGM plates. Animals containing the desired 33bp smo-1(zh156) deletion were identified by PCR amplification using the primers OAF381 (CAC TCG TGT GAG TTG CAT TCT CCA TAG) and OAF383 (CAC GGA AGT GCA CTT CGT TGC TGT C) followed by sequencing. The smo-1(zh156) mutant was back-crossed twice to N2.

### SUMO protein levels by immunoblotting

Worms were collected in H2O supplemented with 20 mM N-ethylmaleimide. For each sample, 50 worms in a volume of 25 ml were mixed with 2x SDS sample buffer, boiled for 5 minutes and stored at -80∼C until analysis. The samples were then reboiled, cooled to RT and treated with DNAse for 10 – 15 minutes. Half of each sample was separated by SDS PAGE and analyzed by immunoblotting using anti-SUMO 6F2. SUMO 6F2 was obtained from the DSHB, where it had been deposited by Pelisch, F. / Hay, R.T. (DSHB Hybridoma Product SUMO 6F2). The immunoblots were quantified with Fiji^109^ and statistical analysis was performed using GraphPad Prism.

### Brood size assay

The brood size assay was performed according to ^110^. The L4 hermaphrodites were selected into individual NGM plates seeded with E. coli strain OP50 and incubated at 20°C. The animals were transferred to a fresh plate every 24h until the end of their self-reproduction period (around fifth day of the adult life). The number of the progeny from each plate was counted. Total brood size is a sum of daily collected data. The experiment was replicated three times, and each experiment included 10 animals of wild-type and the mutant smo-1(zh156) genotype. The hermaphrodites, which were lost or died due to transfer to a new plate, were excluded from the analysis.

### DAPI and SYTO12 staining

L4 hermaphrodites (50-60 animals per genotype) were selected to fresh NGM plate containing OP50 bacteria and incubated at 20°C for 24 h. The adult animals were transferred in M9 buffer to a siliconized Eppendorf tube, spun down for 1min at 3000 rpm, washed with 750 µl of PBS and spun down. Washed animals were incubated in 750 µl ice-cold methanol at -20°C for 5 min and washed twice with 750µl of PBS-T. The animals were incubated in 750 µl DAPI solution (0.1% in PBS) for 15 min and washed twice with 750 µl of PBS-T before mounting in Mowiol. To stain corpses in adult hermaphrodites germline, the SYTO 12 staining protocol was used according to^60^. We scored SYTO 12 positive germ cells per gonad arm of the adult hermaphrodites (29 hours post L4). Three biological replicates of the assay were performed.

### Microscopy and image processing

For Nomarski and fluorescent imaging, live animals were mounted on 4% agarose pads and immobilized by 20 mM tetramisole hydrochloride solution in M9 buffer, unless stated otherwise. To count DAPI-stained nuclei and for SYTO 12 staining, a Leica DM6000 B microscope equipped with Nomarski and fluorescence optics and the Leica Application Suite X software was used. The animals expressing CED-1::GFP were imaged on a Leica DMRA microscope controlled by custom MatLab script and equipped with a beam splitter and two Hamamatsu ORCA-flash4.0L cameras to simultaneously acquire z-stacks in the DIC and GFP channels was used. Images were analyzed and quantified with Fiji software^109^. The L4 hermaphrodites of the strains WS4116 and AH6119 containing the CED-1::GFP reporter were picked to fresh NGM plate seeded with OP50 bacteria and incubated for 24h or 48h at 20°C before being prepared for image analysis. CED-1::GFP-labeled corpses were counted by inspecting the z-stacks recorded across the pachytene zone. DAPI-stained nuclei in the pachytene region were manually counted across z-stacks in fixed and stained animals, and the area corresponding to the pachytene region in each gonad arm was identified through the morphology of the DAPI-stained nuclei.

The statistical analysis was performed by using GraphPad Prism as indicated in the figure legend.

## ACKNOWLEDGEMENTS

We acknowledge Ana Cristina Laranjeira for generating the C. elegans strain AH5891, and thank Klaus "Gerry" Meese for excellent technical support, and former and current lab members for constructs, reagents and helpful discussions. S.M.R. acknowledges the Heidelberg Biosciences International Graduate School (HBIGS). F.J. is grateful for funding from the BIOMS program of Heidelberg University. F.G. acknowledges funding by the Klaus Tschira Foundation. F.M. and F.G. were members of the Cluster CellNetworks.

## AUTHOR CONTRIBUTIONS

S.M.R., A.Fr. and A.Fl. designed and carried out most biochemical experiments, F.J. designed and carried out all Molecular Dynamics simulations, T.R. designed and carried out all NMR experiments, A.Fe. designed and carried out *C. elegans* in vivo experiments. E.M. carried out some biochemical experiments. M.D. carried out some *C. elegans* in vivo experiments. F.G. guided simulations, R.K. guided NMR experiments,

A.H. guided *C.elegans* in vivo experiments, F.M. and A. Fl. guided the overall design of the project. T.R., F.J., S.M.R., A.Fe., A.H., R.K., F.G., A.Fl. and F.M. wrote the manuscript.

## REFERENCES FOR STAR METHODS

Flotho, A., Werner, A., Winter, T., Frank, A.S., Ehret, H. and Melchior, F. (2012). Recombinant reconstitution of sumoylation reactions in vitro. Methods Mol Biol. 832, 93–110.

Hess, B. (2008). P-LINCS: A Parallel Linear Constraint Solver for Molecular Simulation. J Chem Theory Comput. 4, 116–122.

Hutten, S., Flotho, A., Melchior, F. and Kehlenbach, R.H. (2008) The Nup358- RanGAP complex is required for efficient importin alpha/beta-dependent nuclear import. Mol. Biol. Cell 10, 2300–2310.

Johnson, E.S. and Gupta, A.A. (2001) An E3-like factor that promotes SUMO conjugation to the yeast septins. Cell 106, 735–744.

Lindorff-Larsen, K., Piana, S., Palmo, K., Maragakis, P., Klepeis, J.L., Dror, R.O. and Shaw, D.E. (2010). Improved side-chain torsion potentials for the Amber ff99SB protein force field. Proteins 78, 1950–1958.

Mahajan, R., Delphin, C., Guan, T., Gerace, L. and Melchior, F. (1997) A small ubiquitin related polypeptide involved in targeting RanGAP1 to nuclear pore complex protein RanBP2. Cell 88, 97–107.

Melchior, F. (2000). SUMO-1 - Non-Classical Ubiquitin. Annu. Rev. Cell Dev. Biol. 16, 591–626.

Meulmeester, E., Kunze, M., Hsiaoh, H.H., Urlaub, H., Melchior, F. (2008) Mechanism and consequences of paralog specific sumoylation of USP25. Mol. Cell 30, 610–619.

Pettersen E.F., Goddard, T.D., Huang, C.C., Couch, G.S., Greenblatt, D.M., Meng, E.C., Ferrin, T.E. (2004). UCSF Chimera--a visualization system for exploratory research and analysis. J Comput Chem. 25, 1605–1612.

Piana, S., Donchev, A.G., Robustelli, P. and Shaw, D.E. (2015). Water dispersion interactions strongly influence simulated structural properties of disordered protein states. J Phys Chem B., 119, 5113–5123.

Pichler, A., Gast, A., Seeler, J.S., Dejean, A. and Melchior, F. (2002) The nucleoporin RanBP2 is a SUMO1 E3 Ligase. Cell 108, 109–120.

Saitoh, H. and Hinchey, J. (2000). Functional heterogeneity of small ubiquitin-related protein modifiers SUMO-1 versus SUMO-2/3. J Biol Chem 275, 6252–58.

Van Der Spoel, D., Lindahl, E., Hess, B., Groenhof, G., Mark, A.E., Berendsen, H.J. (2005). GROMACS: fast, flexible, and free. J Comput Chem. 26, 1701–1718.

Warnecke, A., Sandalova, T., Achour, A. and Harris, R.A. (2014). PyTMs: a useful PyMOL plugin for modeling common post-translational modifications. BMC Bioinformatics. 15, 370.

Webb, B., and Sali, A. (2014). Comparative Protein Structure Modeling Using MODELLER. Curr Protoc Bioinformatics 47, 5.6.1-32.

Werner, A., Flotho, A. and Melchior, F. (2012). The RanBP2/RanGAP1*SUMO1/Ubc9 Complex Is a Multisubunit SUMO E3 Ligase. Mol Cell 46, 287–298.

## REFERENCES

1. Flotho, A., and Melchior, F. (2013). Sumoylation: a regulatory protein modification in health and disease. Annu Rev Biochem 82, 357–385. 10.1146/annurev-biochem-061909-093311.

2. Gareau, J.R., and Lima, C.D. (2010). The SUMO pathway: emerging mechanisms that shape specificity, conjugation and recognition. Nat Rev Mol Cell Biol 11, 861–871. 10.1038/nrm3011.

3. Melchior, F. (2000). SUMO--nonclassical ubiquitin. Annu Rev Cell Dev Biol 16, 591–626. 10.1146/annurev.cellbio.16.1.591.

4. Vertegaal, A.C.O. (2022). Signalling mechanisms and cellular functions of SUMO. Nat Rev Mol Cell Biol 23, 715–731. 10.1038/s41580-022-00500-y.

5. Golebiowski, F., Matic, I., Tatham, M.H., Cole, C., Yin, Y., Nakamura, A., Cox, J., Barton, G.J., Mann, M., and Hay, R.T. (2009). System-wide changes to SUMO modifications in response to heat shock. Sci Signal 2, ra24. 10.1126/scisignal.2000282.

6. Hendriks, I.A., and Vertegaal, A.C. (2016). A comprehensive compilation of SUMO proteomics. Nat Rev Mol Cell Biol 17, 581–595. 10.1038/nrm.2016.81.

7. Pichler, A., Fatouros, C., Lee, H., and Eisenhardt, N. (2017). SUMO conjugation - a mechanistic view. Biomol Concepts 8, 13–36. 10.1515/bmc-2016-0030.

8. Nayak, A., and Muller, S. (2014). SUMO-specific proteases/isopeptidases: SENPs and beyond. Genome Biol 15, 422. 10.1186/s13059-014-0422-2.

9. Hickey, C.M., Wilson, N.R., and Hochstrasser, M. (2012). Function and regulation of SUMO proteases. Nat Rev Mol Cell Biol 13, 755–766. 10.1038/nrm3478.

10. Novatchkova, M., Budhiraja, R., Coupland, G., Eisenhaber, F., and Bachmair, A. (2004). SUMO conjugation in plants. Planta 220, 1–8. 10.1007/s00425-004-1370-y.

11. Bencsath, K.P., Podgorski, M.S., Pagala, V.R., Slaughter, C.A., and Schulman, B.A. (2002). Identification of a multifunctional binding site on Ubc9p required for Smt3p conjugation. J Biol Chem 277, 47938–47945. 10.1074/jbc.M207442200.

12. Hendriks, I.A., Lyon, D., Young, C., Jensen, L.J., Vertegaal, A.C., and Nielsen, M.L. (2017). Site-specific mapping of the human SUMO proteome reveals co- modification with phosphorylation. Nat Struct Mol Biol 24, 325–336. 10.1038/nsmb.3366.

13. Tatham, M.H., Jaffray, E., Vaughan, O.A., Desterro, J.M., Botting, C.H., Naismith, J.H., and Hay, R.T. (2001). Polymeric chains of SUMO-2 and SUMO-3 are conjugated to protein substrates by SAE1/SAE2 and Ubc9. J Biol Chem 276, 35368–35374. 10.1074/jbc.M104214200.

14. Becker, J., Barysch, S.V., Karaca, S., Dittner, C., Hsiao, H.H., Berriel Diaz, M., Herzig, S., Urlaub, H., and Melchior, F. (2013). Detecting endogenous SUMO targets in mammalian cells and tissues. Nat Struct Mol Biol 20, 525–531. 10.1038/nsmb.2526.

15. Meulmeester, E., Kunze, M., Hsiao, H.H., Urlaub, H., and Melchior, F. (2008). Mechanism and consequences for paralog-specific sumoylation of ubiquitin-specific protease 25. Mol Cell 30, 610–619. 10.1016/j.molcel.2008.03.021.

16. Zhu, J., Zhu, S., Guzzo, C.M., Ellis, N.A., Sung, K.S., Choi, C.Y., and Matunis, M.J. (2008). Small ubiquitin-related modifier (SUMO) binding determines substrate recognition and paralog-selective SUMO modification. J Biol Chem 283, 29405–29415. 10.1074/jbc.M803632200.

17. Pfander, B., Moldovan, G.L., Sacher, M., Hoege, C., and Jentsch, S. (2005). SUMO-modified PCNA recruits Srs2 to prevent recombination during S phase. Nature 436, 428–433. 10.1038/nature03665.

18. Lallemand-Breitenbach, V., Jeanne, M., Benhenda, S., Nasr, R., Lei, M., Peres, L., Zhou, J., Zhu, J., Raught, B., and de The, H. (2008). Arsenic degrades PML or PML- RARalpha through a SUMO-triggered RNF4/ubiquitin-mediated pathway. Nat Cell Biol 10, 547–555. 10.1038/ncb1717.

19. Tatham, M.H., Geoffroy, M.C., Shen, L., Plechanovova, A., Hattersley, N., Jaffray, E.G., Palvimo, J.J., and Hay, R.T. (2008). RNF4 is a poly-SUMO-specific E3 ubiquitin ligase required for arsenic-induced PML degradation. Nat Cell Biol 10, 538-546. 10.1038/ncb1716.

20. Kerscher, O. (2007). SUMO junction-what’s your function? New insights through SUMO-interacting motifs. EMBO Rep 8, 550–555. 10.1038/sj.embor.7400980.

21. Capili, A.D., and Lima, C.D. (2007). Structure and analysis of a complex between SUMO and Ubc9 illustrates features of a conserved E2-Ubl interaction. J Mol Biol 369, 608–618. 10.1016/j.jmb.2007.04.006.

22. Duda, D.M., van Waardenburg, R.C., Borg, L.A., McGarity, S., Nourse, A., Waddell, M.B., Bjornsti, M.A., and Schulman, B.A. (2007). Structure of a SUMO- binding-motif mimic bound to Smt3p-Ubc9p: conservation of a non-covalent ubiquitin- like protein-E2 complex as a platform for selective interactions within a SUMO pathway. J Mol Biol 369, 619–630. 10.1016/j.jmb.2007.04.007.

23. Liu, Q., Jin, C., Liao, X., Shen, Z., Chen, D.J., and Chen, Y. (1999). The binding interface between an E2 (UBC9) and a ubiquitin homologue (UBL1). J Biol Chem 274, 16979–16987. 10.1074/jbc.274.24.16979.

24. Pilla, E., Moller, U., Sauer, G., Mattiroli, F., Melchior, F., and Geiss-Friedlander, R. (2012). A novel SUMO1-specific interacting motif in dipeptidyl peptidase 9 (DPP9) that is important for enzymatic regulation. J Biol Chem 287, 44320–44329. 10.1074/jbc.M112.397224.

25. Diehl, C., Akke, M., Bekker-Jensen, S., Mailand, N., Streicher, W., and Wikstrom, M. (2016). Structural Analysis of a Complex between Small Ubiquitin-like a New Interaction Surface on SUMO. J Biol Chem 291, 12658–12672. 10.1074/jbc.M115.711325.

26. Liu, J., Xue, Z., Zhang, Y., Vann, K.R., Shi, X., and Kutateladze, T.G. (2020). Structural Insight into Binding of the ZZ Domain of HERC2 to Histone H3 and SUMO1. Structure 28, 1225–1230 e1223. 10.1016/j.str.2020.07.003.

27. Vogt, B., and Hofmann, K. (2012). Bioinformatical detection of recognition factors for ubiquitin and SUMO. Methods Mol Biol 832, 249–261. 10.1007/978-1-61779-474-2_18.

28. Sun, H., and Hunter, T. (2012). Poly-small ubiquitin-like modifier (PolySUMO)- binding proteins identified through a string search. J Biol Chem 287, 42071–42083. 10.1074/jbc.M112.410985.

29. Hecker, C.M., Rabiller, M., Haglund, K., Bayer, P., and Dikic, I. (2006). Specification of SUMO1- and SUMO2-interacting motifs. J Biol Chem 281, 16117–16127. 10.1074/jbc.M512757200.

30. Song, J., Durrin, L.K., Wilkinson, T.A., Krontiris, T.G., and Chen, Y. (2004). Identification of a SUMO-binding motif that recognizes SUMO-modified proteins. Proc Natl Acad Sci U S A 101, 14373–14378. 10.1073/pnas.0403498101.

31. Schellenberg, M.J., Lieberman, J.A., Herrero-Ruiz, A., Butler, L.R., Williams, J.G., Munoz-Cabello, A.M., Mueller, G.A., London, R.E., Cortes-Ledesma, F., and Williams, R.S. (2017). ZATT (ZNF451)-mediated resolution of topoisomerase 2 DNA- protein cross-links. Science 357, 1412–1416. 10.1126/science.aam6468.

32. Husnjak, K., and Dikic, I. (2012). Ubiquitin-binding proteins: decoders of ubiquitin-mediated cellular functions. Annu Rev Biochem 81, 291–322. 10.1146/annurev-biochem-051810-094654.

33. Stehmeier, P., and Muller, S. (2009). Phospho-regulated SUMO interaction modules connect the SUMO system to CK2 signaling. Mol Cell 33, 400–409. 10.1016/j.molcel.2009.01.013.

34. Cappadocia, L., Mascle, X.H., Bourdeau, V., Tremblay-Belzile, S., Chaker- Margot, M., Lussier-Price, M., Wada, J., Sakaguchi, K., Aubry, M., Ferbeyre, G., and Omichinski, J.G. (2015). Structural and functional characterization of the phosphorylation-dependent interaction between PML and SUMO1. Structure 23, 126-138. 10.1016/j.str.2014.10.015.

35. Chang, C.C., Naik, M.T., Huang, Y.S., Jeng, J.C., Liao, P.H., Kuo, H.Y., Ho, C.C., Hsieh, Y.L., Lin, C.H., Huang, N.J., et al. (2011). Structural and functional roles of Daxx SIM phosphorylation in SUMO paralog-selective binding and apoptosis modulation. Mol Cell 42, 62–74. 10.1016/j.molcel.2011.02.022.

36. Escobar-Cabrera, E., Okon, M., Lau, D.K., Dart, C.F., Bonvin, A.M., and McIntosh, L.P. (2011). Characterizing the N- and C-terminal Small ubiquitin-like modifier (SUMO)-interacting motifs of the scaffold protein DAXX. J Biol Chem 286, 19816–19829. 10.1074/jbc.M111.231647.

37. Namanja, A.T., Li, Y.J., Su, Y., Wong, S., Lu, J., Colson, L.T., Wu, C., Li, S.S., and Chen, Y. (2012). Insights into high affinity small ubiquitin-like modifier (SUMO) recognition by SUMO-interacting motifs (SIMs) revealed by a combination of NMR and peptide array analysis. J Biol Chem 287, 3231–3240. 10.1074/jbc.M111.293118.

38. Sekiyama, N., Ikegami, T., Yamane, T., Ikeguchi, M., Uchimura, Y., Baba, D., Ariyoshi, M., Tochio, H., Saitoh, H., and Shirakawa, M. (2008). Structure of the small ubiquitin-like modifier (SUMO)-interacting motif of MBD1-containing chromatin- associated factor 1 bound to SUMO-3. J Biol Chem 283, 35966–35975. 10.1074/jbc.M802528200.

39. Bayer, P., Arndt, A., Metzger, S., Mahajan, R., Melchior, F., Jaenicke, R., and Becker, J. (1998). Structure determination of the small ubiquitin-related modifier SUMO-1. J Mol Biol 280, 275–286. 10.1006/jmbi.1998.1839.

40. Bylebyl, G.R., Belichenko, I., and Johnson, E.S. (2003). The SUMO isopeptidase Ulp2 prevents accumulation of SUMO chains in yeast. J Biol Chem 278, 44113–44120. 10.1074/jbc.M308357200.

41. Newman, H.A., Meluh, P.B., Lu, J., Vidal, J., Carson, C., Lagesse, E., Gray, J.J., Boeke, J.D., and Matunis, M.J. (2017). A high throughput mutagenic analysis of yeast sumo structure and function. PLoS Genet 13, e1006612. 10.1371/journal.pgen.1006612.

42. Lussier-Price, M., Wahba, H.M., Mascle, X.H., Cappadocia, L., Bourdeau, V., Gagnon, C., Igelmann, S., Sakaguchi, K., Ferbeyre, G., and Omichinski, J.G. (2022). Zinc controls PML nuclear body formation through regulation of a paralog specific auto- inhibition in SUMO1. Nucleic Acids Res 50, 8331–8348. 10.1093/nar/gkac620.

43. Srikumar, T., Lewicki, M.C., Costanzo, M., Tkach, J.M., van Bakel, H., Tsui, K., Johnson, E.S., Brown, G.W., Andrews, B.J., Boone, C., et al. (2013). Global analysis of SUMO chain function reveals multiple roles in chromatin regulation. J Cell Biol 201, 145–163. 10.1083/jcb.201210019.

44. Pichler, A., Gast, A., Seeler, J.S., Dejean, A., and Melchior, F. (2002). The nucleoporin RanBP2 has SUMO1 E3 ligase activity. Cell 108, 109–120.

45. Tatham, M.H., Kim, S., Jaffray, E., Song, J., Chen, Y., and Hay, R.T. (2005). Unique binding interactions among Ubc9, SUMO and RanBP2 reveal a mechanism for SUMO paralog selection. Nat Struct Mol Biol 12, 67–74. 10.1038/nsmb878.

46. Knipscheer, P., van Dijk, W.J., Olsen, J.V., Mann, M., and Sixma, T.K. (2007). Noncovalent interaction between Ubc9 and SUMO promotes SUMO chain formation. EMBO J 26, 2797–2807. 10.1038/sj.emboj.7601711.

47. Takahashi, Y., Iwase, M., Konishi, M., Tanaka, M., Toh-e, A., and Kikuchi, Y. (1999). Smt3, a SUMO-1 homolog, is conjugated to Cdc3, a component of septin rings at the mother-bud neck in budding yeast. Biochem Biophys Res Commun 259, 582-587. 10.1006/bbrc.1999.0821.

48. Vertegaal, A.C. (2010). SUMO chains: polymeric signals. Biochem Soc Trans 38, 46–49. 10.1042/BST0380046.

49. Takahashi, Y., and Kikuchi, Y. (2005). Yeast PIAS-type Ull1/Siz1 is composed of SUMO ligase and regulatory domains. J Biol Chem 280, 35822–35828. 10.1074/jbc.M506794200.

50. Pelisch, F., Tammsalu, T., Wang, B., Jaffray, E.G., Gartner, A., and Hay, R.T. (2017). A SUMO-Dependent Protein Network Regulates Chromosome Congression during Oocyte Meiosis. Mol Cell 65, 66–77. 10.1016/j.molcel.2016.11.001.

51. Wand, A.J., Urbauer, J.L., McEvoy, R.P., and Bieber, R.J. (1996). Internal dynamics of human ubiquitin revealed by 13C-relaxation studies of randomly fractionally labeled protein. Biochemistry 35, 6116–6125. 10.1021/bi9530144.

52. Borgia, A., Borgia, M.B., Bugge, K., Kissling, V.M., Heidarsson, P.O., Fernandes, C.B., Sottini, A., Soranno, A., Buholzer, K.J., Nettels, D., et al. (2018). Extreme disorder in an ultrahigh-affinity protein complex. Nature 555, 61–66. 10.1038/nature25762.

53. Sharma, R., Raduly, Z., Miskei, M., and Fuxreiter, M. (2015). Fuzzy complexes: Specific binding without complete folding. FEBS Lett 589, 2533–2542. 10.1016/j.febslet.2015.07.022.

54. Jones, D.T. (1999). Protein secondary structure prediction based on position-specific scoring matrices. J Mol Biol 292, 195–202. 10.1006/jmbi.1999.3091.

55. Ahmed, M.C., Papaleo, E., and Lindorff-Larsen, K. (2018). How well do force fields capture the strength of salt bridges in proteins? Peerj 6. ARTN e4967 10.7717/peerj.4967.

56. Matic, I., Macek, B., Hilger, M., Walther, T.C., and Mann, M. (2008). Phosphorylation of SUMO-1 occurs in vivo and is conserved through evolution. J Proteome Res 7, 4050–4057. 10.1021/pr800368m.

57. Ullmann, R., Chien, C.D., Avantaggiati, M.L., and Muller, S. (2012). An acetylation switch regulates SUMO-dependent protein interaction networks. Mol Cell 46, 759–770. 10.1016/j.molcel.2012.04.006.

58. Pelisch, F., Sonneville, R., Pourkarimi, E., Agostinho, A., Blow, J.J., Gartner, A., and Hay, R.T. (2014). Dynamic SUMO modification regulates mitotic chromosome assembly and cell cycle progression in Caenorhabditis elegans. Nat Commun 5, 5485. 10.1038/ncomms6485.

59. Reichman, R., Shi, Z., Malone, R., and Smolikove, S. (2018). Mitotic and Meiotic Functions for the SUMOylation Pathway in the Caenorhabditis elegans Germline. Genetics 208, 1421–1441. 10.1534/genetics.118.300787.

60. Gumienny, T.L., Lambie, E., Hartwieg, E., Horvitz, H.R., and Hengartner, M.O. (1999). Genetic control of programmed cell death in the Caenorhabditis elegans hermaphrodite germline. Development 126, 1011–1022. 10.1242/dev.126.5.1011.

61. Schumacher, B., Hanazawa, M., Lee, M.H., Nayak, S., Volkmann, K., Hofmann, E.R., Hengartner, M., Schedl, T., and Gartner, A. (2005). Translational repression of C. elegans p53 by GLD-1 regulates DNA damage-induced apoptosis. Cell 120, 357–368. 10.1016/j.cell.2004.12.009.

62. Ward, J.J., Sodhi, J.S., McGuffin, L.J., Buxton, B.F., and Jones, D.T. (2004). Prediction and functional analysis of native disorder in proteins from the three kingdoms of life. J Mol Biol 337, 635–645. 10.1016/j.jmb.2004.02.002.

63. Dunker, A.K., Brown, C.J., Lawson, J.D., Iakoucheva, L.M., and Obradovic, Z. (2002). Intrinsic disorder and protein function. Biochemistry 41, 6573–6582.

64. Uversky, V.N. (2002). Natively unfolded proteins: a point where biology waits for physics. Protein Sci 11, 739–756. 10.1110/ps.4210102.

65. Oldfield, C.J., and Dunker, A.K. (2014). Intrinsically disordered proteins and intrinsically disordered protein regions. Annu Rev Biochem 83, 553–584. 10.1146/annurev-biochem-072711-164947.

66. Tompa, P. (2002). Intrinsically unstructured proteins. Trends Biochem Sci 27, 527–533.

67. Ulrich, H.D. (2008). The fast-growing business of SUMO chains. Mol Cell 32, 301–305. 10.1016/j.molcel.2008.10.010.

68. Gartner, A., Wagner, K., Holper, S., Kunz, K., Rodriguez, M.S., and Muller, S. (2018). Acetylation of SUMO2 at lysine 11 favors the formation of non-canonical SUMO chains. EMBO Rep 19. 10.15252/embr.201846117.

69. Grana-Montes, R., Marinelli, P., Reverter, D., and Ventura, S. (2014). N-terminal protein tails act as aggregation protective entropic bristles: the SUMO case. Biomacromolecules 15, 1194–1203. 10.1021/bm401776z.

70. Gianni, S., Dogan, J., and Jemth, P. (2016). Coupled binding and folding of intrinsically disordered proteins: what can we learn from kinetics? Curr Opin Struct Biol 36, 18–24. 10.1016/j.sbi.2015.11.012.

71. Lacy, E.R., Filippov, I., Lewis, W.S., Otieno, S., Xiao, L., Weiss, S., Hengst, L., and Kriwacki, R.W. (2004). p27 binds cyclin-CDK complexes through a sequential mechanism involving binding-induced protein folding. Nat Struct Mol Biol 11, 358–364. 10.1038/nsmb746.

72. Tompa, P., and Fuxreiter, M. (2008). Fuzzy complexes: polymorphism and structural disorder in protein-protein interactions. Trends Biochem Sci 33, 2–8. 10.1016/j.tibs.2007.10.003.

73. Brzovic, P.S., Heikaus, C.C., Kisselev, L., Vernon, R., Herbig, E., Pacheco, D., Warfield, L., Littlefield, P., Baker, D., Klevit, R.E., and Hahn, S. (2011). The acidic transcription activator Gcn4 binds the mediator subunit Gal11/Med15 using a simple protein interface forming a fuzzy complex. Mol Cell 44, 942–953. 10.1016/j.molcel.2011.11.008.

74. Milles, S., Mercadante, D., Aramburu, I.V., Jensen, M.R., Banterle, N., Koehler, C., Tyagi, S., Clarke, J., Shammas, S.L., Blackledge, M., et al. (2015). Plasticity of an ultrafast interaction between nucleoporins and nuclear transport receptors. Cell 163, 734–745. 10.1016/j.cell.2015.09.047.

75. Arbesu, M., Maffei, M., Cordeiro, T.N., Teixeira, J.M., Perez, Y., Bernado, P., Roche, S., and Pons, M. (2017). The Unique Domain Forms a Fuzzy Intramolecular Complex in Src Family Kinases. Structure 25, 630–640 e634. 10.1016/j.str.2017.02.011.

76. Lin, Y.H., Qiu, D.C., Chang, W.H., Yeh, Y.Q., Jeng, U.S., Liu, F.T., and Huang, J.R. (2017). The intrinsically disordered N-terminal domain of galectin-3 dynamically mediates multisite self-association of the protein through fuzzy interactions. J Biol Chem 292, 17845–17856. 10.1074/jbc.M117.802793.

77. Jin, F., Yu, C., Lai, L., and Liu, Z. (2013). Ligand clouds around protein clouds: a scenario of ligand binding with intrinsically disordered proteins. PLoS Comput Biol 9, e1003249. 10.1371/journal.pcbi.1003249.

78. Pufall, M.A., and Graves, B.J. (2002). Autoinhibitory domains: modular effectors of cellular regulation. Annu Rev Cell Dev Biol 18, 421–462. 10.1146/annurev.cellbio.18.031502.133614.

79. Wright, P.E., and Dyson, H.J. (2015). Intrinsically disordered proteins in cellular signalling and regulation. Nat Rev Mol Cell Biol 16, 18–29. 10.1038/nrm3920.

80. Mascle, X.H., Gagnon, C., Wahba, H.M., Lussier-Price, M., Cappadocia, L., Sakaguchi, K., and Omichinski, J.G. (2019). Acetylation of SUMO1 Alters Interactions with the SIMs of PML and Daxx in a Protein-Specific Manner. Structure. 10.1016/j.str.2019.11.019.

81. Zilio, N., Eifler-Olivi, K., and Ulrich, H.D. (2017). Functions of SUMO in the Maintenance of Genome Stability. Adv Exp Med Biol 963, 51–87. 10.1007/978-3-319-50044-7_4.

82. Hutten, S., Flotho, A., Melchior, F., and Kehlenbach, R.H. (2008). The Nup358- RanGAP complex is required for efficient importin alpha/beta-dependent nuclear import. Mol Biol Cell 19, 2300–2310. 10.1091/mbc.E07-12-1279.

83. Zhou, Z., Hartwieg, E., and Horvitz, H.R. (2001). CED-1 is a transmembrane receptor that mediates cell corpse engulfment in C-elegans. Cell 104, 43–56. Doi 10.1016/S0092-8674(01)00190-8.

84. Barstead, R., Moulder, G., Cobb, B., Frazee, S., Henthorn, D., Holmes, J., Jerebie, D., Landsdale, M., Osborn, J., Pritchett, C., et al. (2012). Large-Scale Screening for Targeted Knockouts in the Caenorhabditis elegans Genome. G3-Genes Genom Genet 2, 1415–1425. 10.1534/g3.112.003830.

85. Werner, A., Flotho, A., and Melchior, F. (2012). The RanBP2/RanGAP1*SUMO1/Ubc9 complex is a multisubunit SUMO E3 ligase. Mol Cell 46, 287–298. 10.1016/j.molcel.2012.02.017.

86. Johnson, E.S., and Gupta, A.A. (2001). An E3-like factor that promotes SUMO conjugation to the yeast septins. Cell 106, 735–744. 10.1016/s0092-8674(01)00491-3.

87. Warnecke, A., Sandalova, T., Achour, A., and Harris, R.A. (2014). PyTMs: a useful PyMOL plugin for modeling common post-translational modifications. BMC Bioinformatics 15, 370. 10.1186/s12859-014-0370-6.

88. Van der Spoel, D., Lindahl, E., Hess, B., Groenhof, G., Mark, A.E., and Berendsen, H.J.C. (2005). GROMACS: Fast, flexible, and free. Journal of Computational Chemistry 26, 1701–1718. 10.1002/jcc.20291.

89. Pettersen, E.F., Goddard, T.D., Huang, C.C., Couch, G.S., Greenblatt, D.M., Meng, E.C., and Ferrin, T.E. (2004). UCSF Chimera--a visualization system for exploratory research and analysis. J Comput Chem 25, 1605–1612. 10.1002/jcc.20084.

90. Webb, B., and Sali, A. (2014). Comparative Protein Structure Modeling Using MODELLER. Curr Protoc Bioinformatics 47, 5 6 1-32. 10.1002/0471250953.bi0506s47.

91. Saitoh, H., and Hinchey, J. (2000). Functional heterogeneity of small ubiquitin-related protein modifiers SUMO-1 versus SUMO-2/3. J Biol Chem 275, 6252–6258. 10.1074/jbc.275.9.6252.

92. Flotho, A., Werner, A., Winter, T., Frank, A.S., Ehret, H., and Melchior, F. (2012). Recombinant reconstitution of sumoylation reactions in vitro. Methods Mol Biol 832, 93–110. 10.1007/978-1-61779-474-2_5.

93. Mahajan, R., Delphin, C., Guan, T., Gerace, L., and Melchior, F. (1997). A small ubiquitin-related polypeptide involved in targeting RanGAP1 to nuclear pore complex protein RanBP2. Cell 88, 97–107.

94. Schneider, C.A., Rasband, W.S., and Eliceiri, K.W. (2012). NIH Image to ImageJ: 25 years of image analysis. Nat Methods 9, 671–675. 10.1038/nmeth.2089.

95. Delaglio, F., Grzesiek, S., Vuister, G.W., Zhu, G., Pfeifer, J., and Bax, A. (1995). NMRPipe: a multidimensional spectral processing system based on UNIX pipes. J Biomol NMR 6, 277–293. 10.1007/BF00197809.

96. Johnson, B.A., and Blevins, R.A. (1994). NMR View: A computer program for the visualization and analysis of NMR data. J Biomol NMR 4, 603–614. 10.1007/BF00404272.

97. Naik, M.T., Chang, C.C., Naik, N.M., Kung, C.C., Shih, H.M., and Huang, T.H. (2011). NMR chemical shift assignments of a complex between SUMO-1 and SIM peptide derived from the C-terminus of Daxx. Biomol NMR Assign 5, 75–77. 10.1007/s12104-010-9271-4.

98. Mossessova, E., and Lima, C.D. (2000). Ulp1-SUMO crystal structure and genetic analysis reveal conserved interactions and a regulatory element essential for cell growth in yeast. Mol Cell 5, 865–876. 10.1016/s1097-2765(00)80326-3.

99. Surana, P., Gowda, C.M., Tripathi, V., Broday, L., and Das, R. (2017). Structural and functional analysis of SMO-1, the SUMO homolog in Caenorhabditis elegans. PLoS One 12, e0186622. 10.1371/journal.pone.0186622.

100. Schrodinger, LLC (2015). The PyMOL Molecular Graphics System, Version 1.8.

101. Papamokos, G.V., Tziatzos, G., Papageorgiou, D.G., Georgatos, S.D., Politou, A.S., and Kaxiras, E. (2012). Structural role of RKS motifs in chromatin interactions: a molecular dynamics study of HP1 bound to a variably modified histone tail. Biophys J 102, 1926–1933. 10.1016/j.bpj.2012.03.030.

102. Homeyer, N., Horn, A.H., Lanig, H., and Sticht, H. (2006). AMBER force-field parameters for phosphorylated amino acids in different protonation states: phosphoserine, phosphothreonine, phosphotyrosine, and phosphohistidine. J Mol Model 12, 281–289. 10.1007/s00894-005-0028-4.

103. Lindorff-Larsen, K., Piana, S., Palmo, K., Maragakis, P., Klepeis, J.L., Dror, R.O., and Shaw, D.E. (2010). Improved side-chain torsion potentials for the Amber ff99SB protein force field. Proteins 78, 1950–1958. 10.1002/prot.22711.

104. Piana, S., Donchev, A.G., Robustelli, P., and Shaw, D.E. (2015). Water dispersion interactions strongly influence simulated structural properties of disordered protein states. J Phys Chem B 119, 5113–5123. 10.1021/jp508971m.

105. Hess, B. (2008). P-LINCS: A Parallel Linear Constraint Solver for Molecular Simulation. J Chem Theory Comput 4, 116–122. 10.1021/ct700200b.

106. Han, B., Liu, Y., Ginzinger, S.W., and Wishart, D.S. (2011). SHIFTX2: significantly improved protein chemical shift prediction. J Biomol NMR 50, 43–57. 10.1007/s10858-011-9478-4.

107. Brenner, S. (1974). The genetics of Caenorhabditis elegans. Genetics 77, 71–94. 10.1093/genetics/77.1.71.

108. Arribere, J.A., Bell, R.T., Fu, B.X., Artiles, K.L., Hartman, P.S., and Fire, A.Z. (2014). Efficient marker-free recovery of custom genetic modifications with CRISPR/Cas9 in Caenorhabditis elegans. Genetics 198, 837–846. 10.1534/genetics.114.169730.

109. Schindelin, J., Arganda-Carreras, I., Frise, E., Kaynig, V., Longair, M., Pietzsch, T., Preibisch, S., Rueden, C., Saalfeld, S., Schmid, B., et al. (2012). Fiji: an open-source platform for biological-image analysis. Nat Methods *9*, 676-682. 10.1038/nmeth.2019.

110. Chawla, D.G., Shah, R.V., Barth, Z.K., Lee, J.D., Badecker, K.E., Naik, A., Brewster, M.M., Salmon, T.P., and Peel, N. (2016). Caenorhabditis elegans glutamylating enzymes function redundantly in male mating. Biol Open 5, 1290–1298. 10.1242/bio.017442.

