## Supplemental Figures for "The intrinsically disordered N-terminus of SUMO1 is an intramolecular inhibitor of SUMO1 interactions"

### Figure S1

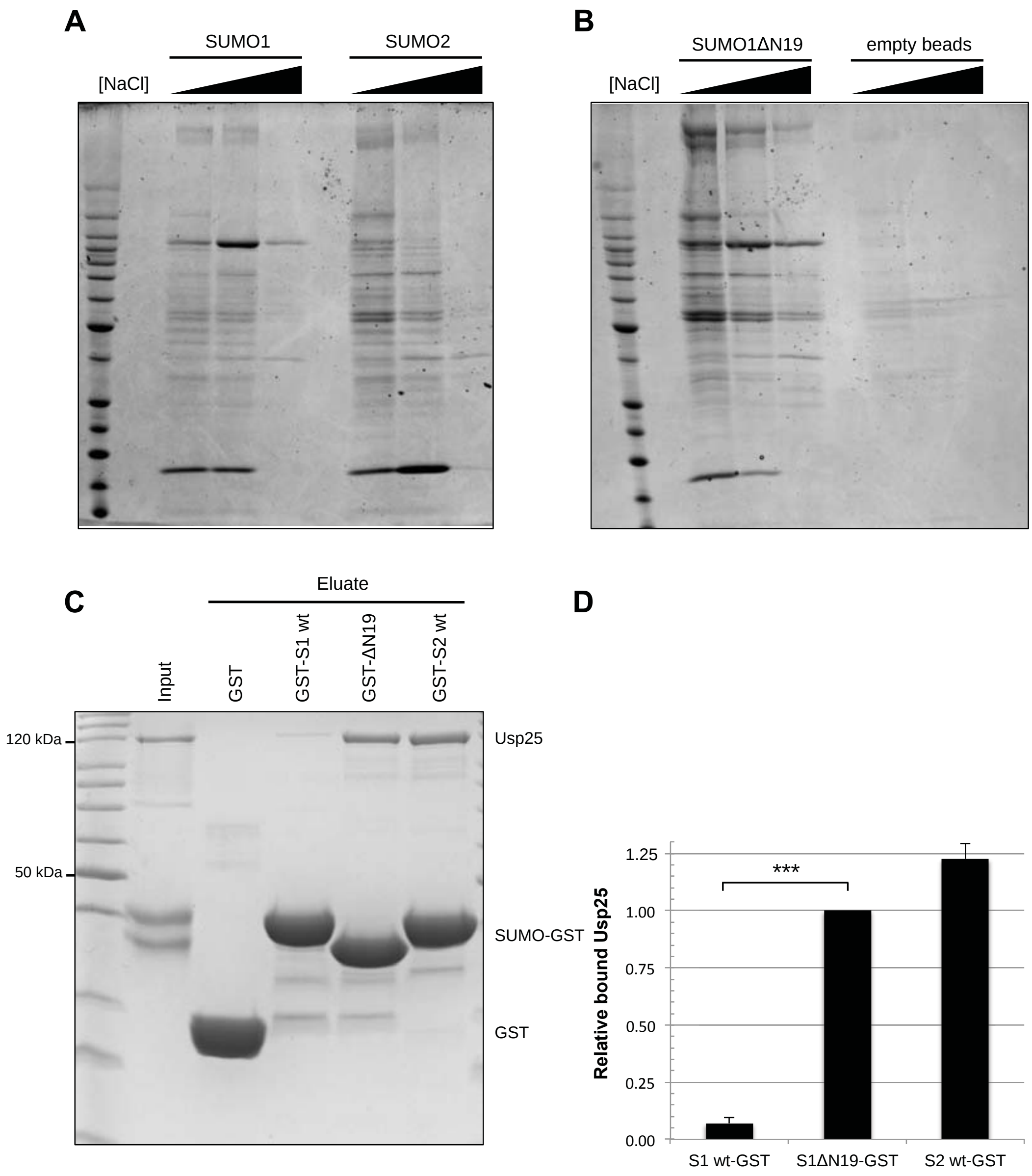
A
SUMO1
SUMO2
[NaCl]
B
SUMO1ΔN19
empty beads
[NaCl]
C
Eluate
Input
GST
GST-S1 wt
GST-ΔN19
GST-S2 wt
120 kDa
50 kDa
Usp25
SUMO-GST
GST
D
Relative bound Usp25
1.25
1.00
0.75
0.50
0.25
0.00
***
S1 wt-GST
S1ΔN19-GST
S2 wt-GST

**Figure S1 Pull-down experiments with SUMO1 and 2, related to Figure 1.**

(A) and (B) SUMO1, SUMO2 and SUMO1ΔN19 were immobilized on sepharose beads and incubated with HeLa cell lysates. Interacting proteins were eluted with increasing salt concentrations and analyzed by SDS-PAGE followed by Coomassie staining. Beads without immobilized protein on it were used as a negative control – see (B). As mentioned in the main text, Fig. S1A is identical to Fig. 1A in (Pilla et al., 2012) and is thus not novel, yet is shown here again to illustrate the starting observation for this paper; Fig. S1B is part of the work for that paper, but had not been published yet.

(C) 10 µg of recombinant Usp25 were incubated with glutathione beads loaded with SUMO1-GST, SUMO1ΔN19-GST or SUMO2-GST for 1 h at 4 °C, washed three times and eluted using 2x SDS-Sample buffer. 5 % of the inputs and 100 % of the eluates were analyzed by SDS-PAGE followed by Coomassie staining. (D) The amounts of Usp25 in the eluates of SUMO pull-downs in (C) were quantified relative to the GST-tagged SUMO proteins using ImageJ. Data were analyzed using a paired Student's t-test ($n = 3$). Protein amounts in the eluates are depicted relative to the pull-down with SUMO1ΔN19-GST. Error bars represent one standard deviation. Asterisks indicate p-values: ***: $P \leq 0.001$.

**Table S1. Overview of proteins (and their variants) for which MD simulations were performed in this work, including the residue borders of the respective N-termini, cores and the SIM-binding regions; each simulation was done with 8 trajectories and a simulation time per trajectory of 1 μs, related to Figures 3, 5 and 6.**

| Protein | N-terminus | Core | SIM-binding groove | Variants |
|---|---|---|---|---|
| human SUMO1 | 1-18 | 22-92 | 32-40, 44-54 | wt<br>ED11,12KK<br>pS2<br>pS9<br>pS9pT10 |
| human SUMO2 | 1-12 | 16-85 | 27-35, 40-50 | wt<br>AcK7<br>pT12<br>AcK11pT12 |
| yeast Smt3 | 1-20 | 24-92 | 33-41, 45-55 | wt |
| worm Smo-1 | 1-11 | 15-84 | 24-32, 37-47 | wt<br>DD3,4KK |

**Figure S2**

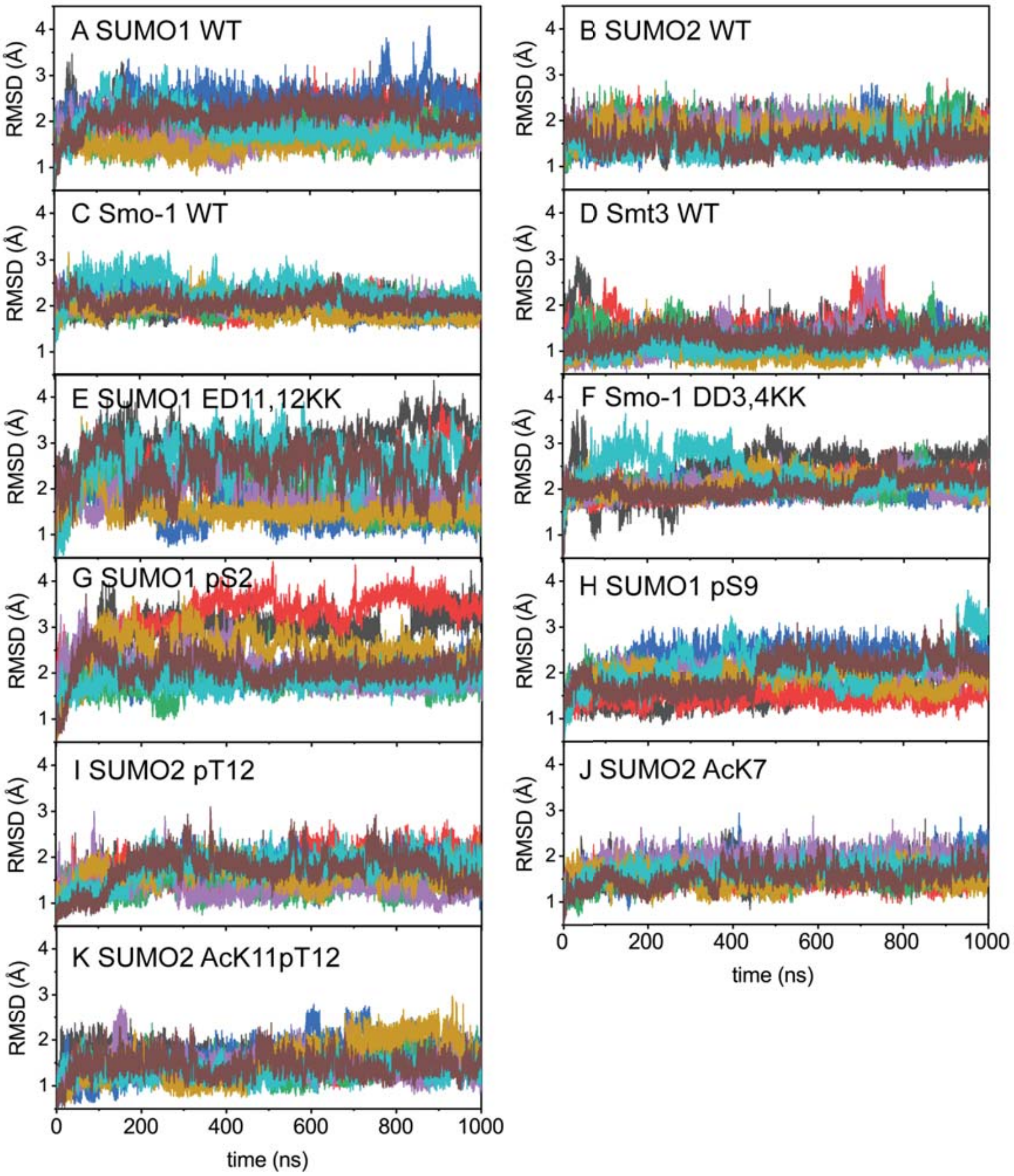
A SUMO1 WT
B SUMO2 WT
C Smo-1 WT
D Smt3 WT
E SUMO1 ED11,12KK
F Smo-1 DD3,4KK
G SUMO1 pS2
H SUMO1 pS9
I SUMO2 pT12
J SUMO2 AcK7
K SUMO2 AcK11pT12
RMSD (Å)
time (ns)
0
200
400
600
800
1000
1
2
3
4

**Figure S2 Structural stability of SUMOs and their variants during MD simulations, related to Figures 3, 5 and 6**

Root-mean-square deviation (RMSD) against the initial structures for Cα atoms of the core of SUMO1, SUMO2, *C. elegans* Smo-1 and yeast Smt3 in their wild type (A) - (D), indicated mutants (E) - (F) and/or post-translationally modified forms (G) - (K). See Table S2 for the definition of residues of the N-terminus in each protein. Each panel shows the RMSD over independent MD simulations (shown in different colors).

All the MD simulations for different proteins/protein variants show relatively stable RMSD values with fluctuations at an average of 2-3 Å; despite the binding/unbinding behavior of the N-terminus, the SUMO cores remain folded and stable during MD simulations.

**Figure S3**

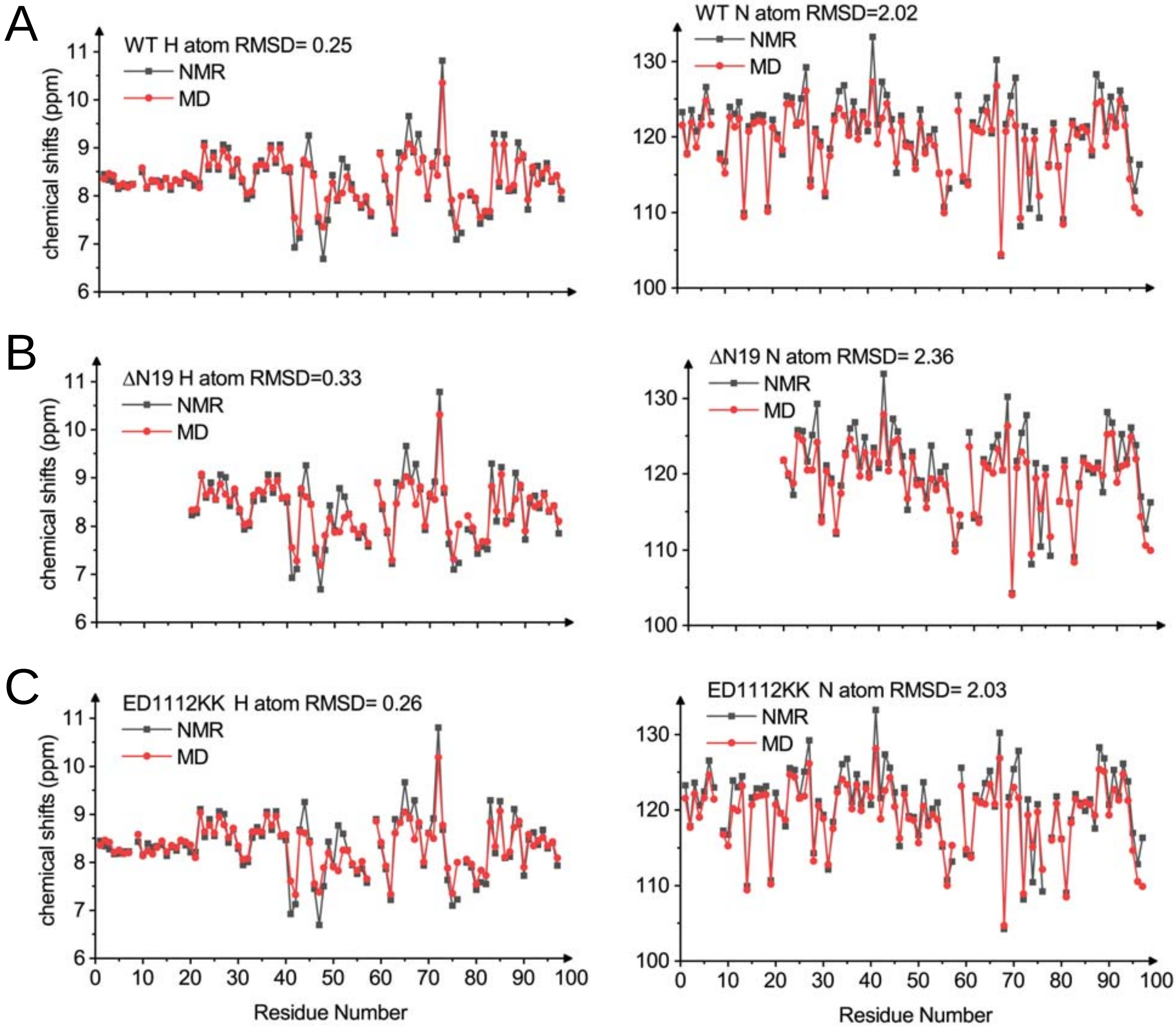
A
WT H atom RMSD= 0.25
NMR
MD
chemical shifts (ppm)
WT N atom RMSD=2.02
B
ΔN19 H atom RMSD=0.33
ΔN19 N atom RMSD= 2.36
C
ED1112KK H atom RMSD= 0.26
ED1112KK N atom RMSD= 2.03
Residue Number

**Figure S3 Comparisons of experimental and theoretical chemical shifts, related to Fig. 3 and 5.**

Chemical shifts obtained by NMR experiments (black) and shifts back-calculated from MD simulations (red) were plotted for SUMO1 wt (A), ΔN19 (B) and ED11,12KK (C). Plots in the left column show shifts in the H dimension, those in the right column shift in the N dimension. For each plot, the RMSD was calculated.

**Figure S4**

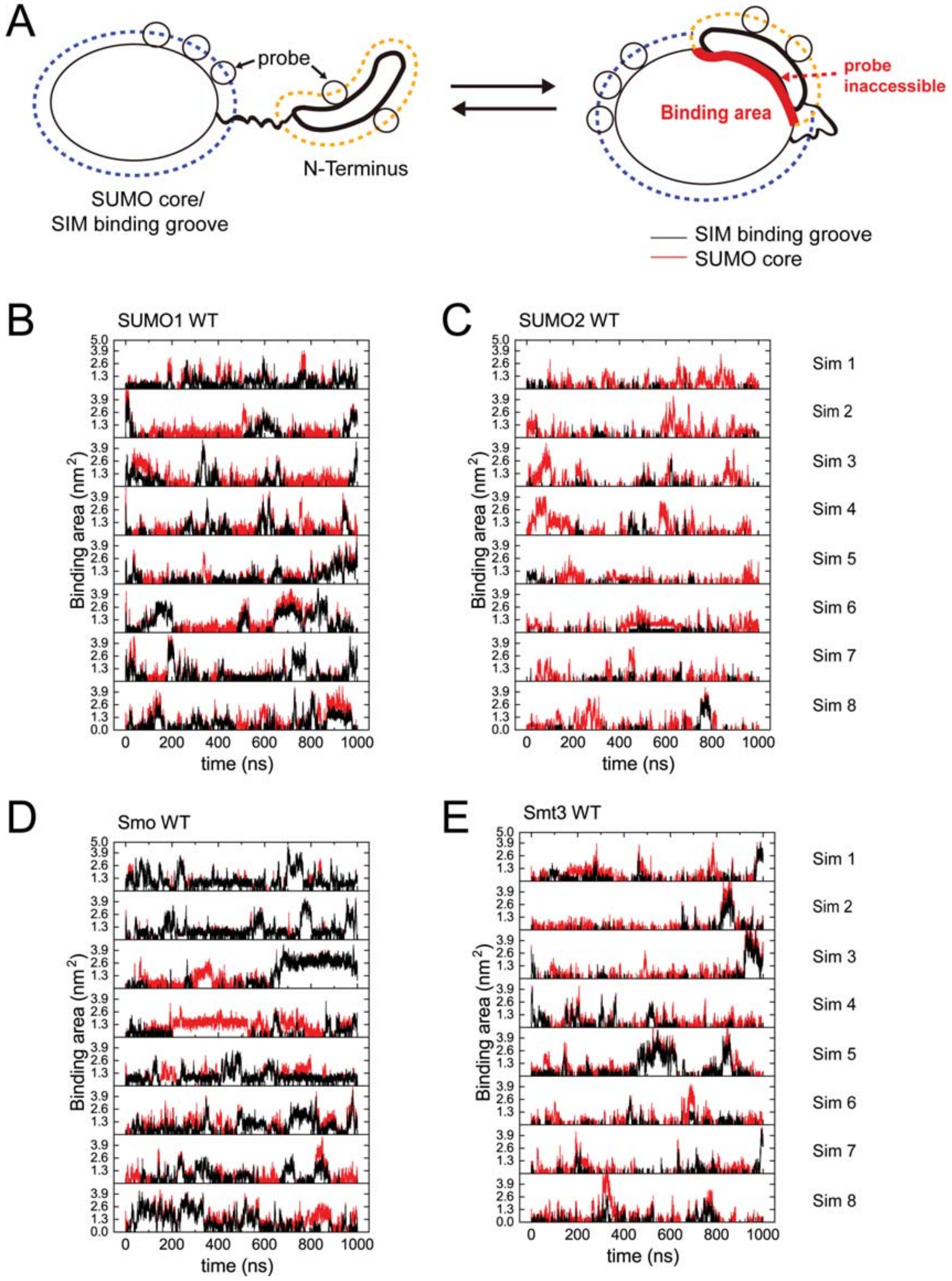
A
probe
N-Terminus
SUMO core/
SIM binding groove
probe
inaccessible
Binding area
SIM binding groove
SUMO core
B
SUMO1 WT
C
SUMO2 WT
Binding area (nm$^2$)
time (ns)
Sim 1
Sim 2
Sim 3
Sim 4
Sim 5
Sim 6
Sim 7
Sim 8
D
Smo WT
E
Smt3 WT

**Figure S4 Quantification of the binding area between the N-terminus and SIM binding groove of the SUMO core during MD simulations, related to Figures 3.**
(A) The binding and unbinding process of the N-terminus (orange dashed line) and the SUMO core/SIM-binding groove (blue dashed line) was quantified as the change of the solvent-accessible surface area (SASA), measured as the area accessible to a spherical probe (black circles, size of water molecule): Upon binding of the N-terminus to the SUMO core/SIM binding groove, the binding sites would be inaccessible for the probe (red) and thus representing the binding area. (B) - (E) Time evolution of the binding area of the N-terminus with the SUMO core (red) and the SIM-binding groove (black) during MD simulations for wild-type SUMO1, SUMO2, Smo-1 and Smt3. See Table S2 for the definition of residues of the N-terminus, core and SIM-binding groove in each protein, respectively. The respective last three residues of the N-terminus adjacent to the SUMO core were ignored to leave about 2 nm distance between the N-terminus and the SUMO core and SIM-binding groove, respectively.

**Figure S5**

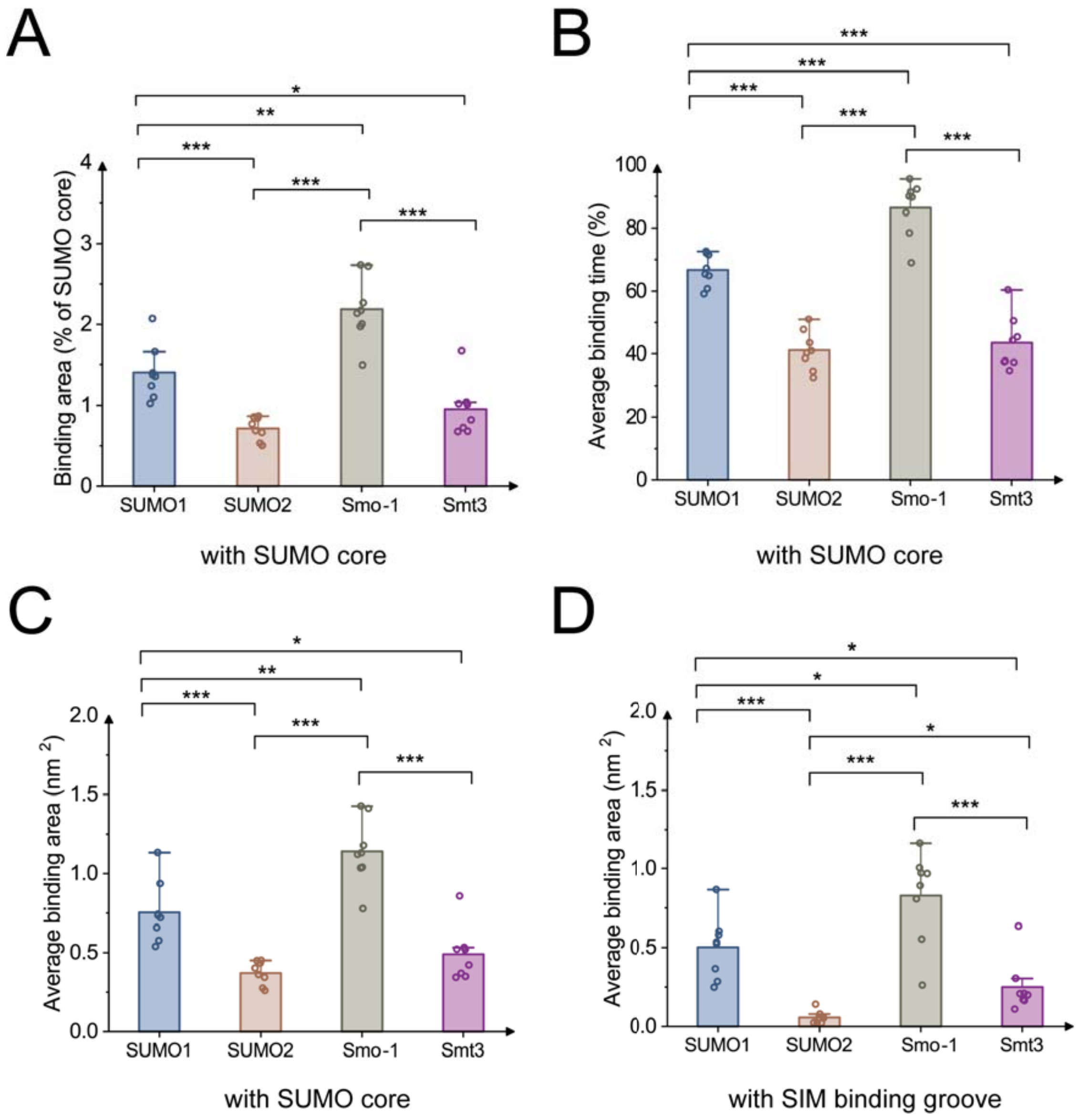

A
*
**
***
***
***
Binding area (% of SUMO core)
0
1
2
3
4
SUMO1
SUMO2
Smo-1
Smt3
with SUMO core
B
***
***
***
***
***
Average binding time (%)
0
20
40
60
80
100
SUMO1
SUMO2
Smo-1
Smt3
with SUMO core
C
*
**
***
***
***
Average binding area (nm 2)
0.0
0.5
1.0
1.5
2.0
SUMO1
SUMO2
Smo-1
Smt3
with SUMO core
D
*
*
***
*
***
***
Average binding area (nm 2)
0.0
0.5
1.0
1.5
2.0
SUMO1
SUMO2
Smo-1
Smt3
with SIM binding groove

### Figure S5 - continued

E

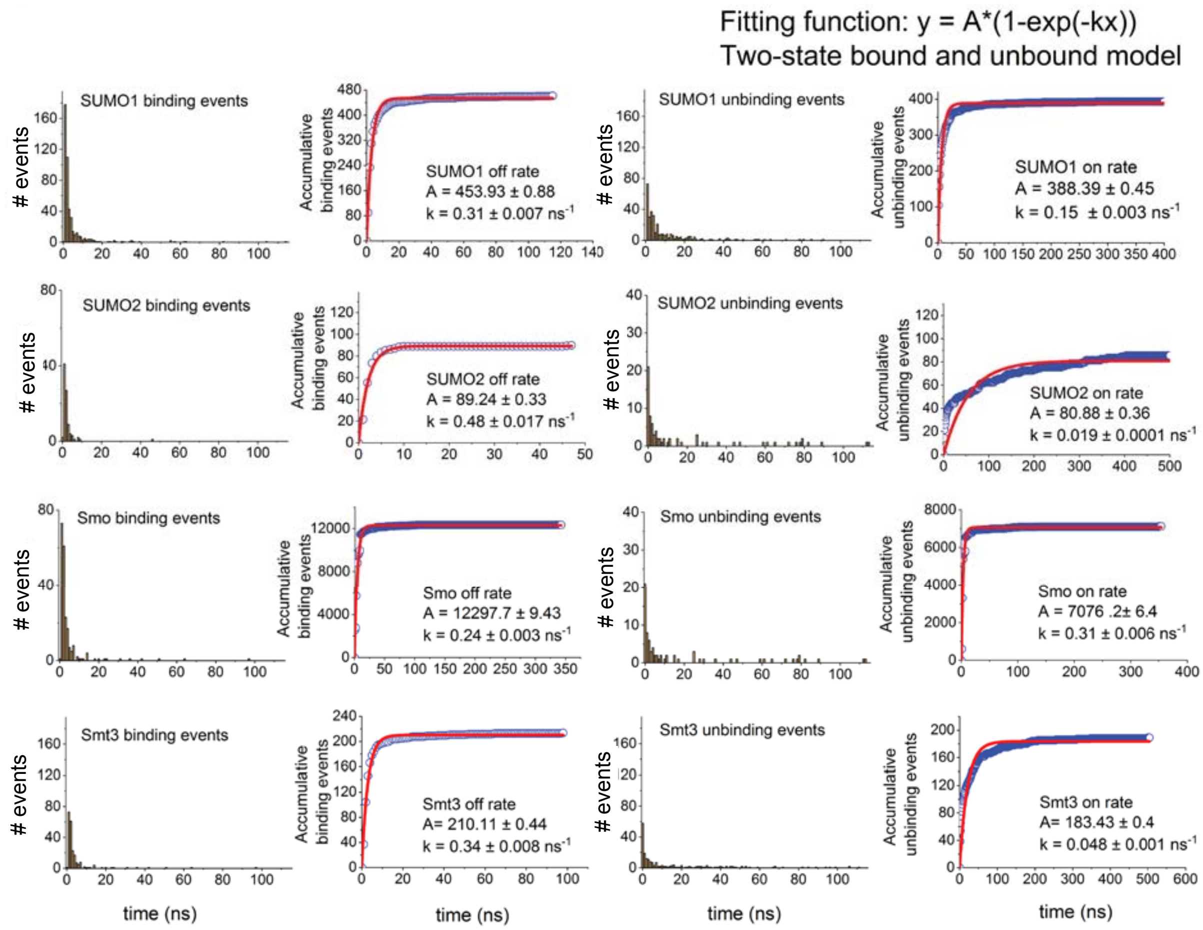

**Figure S5 MD simulations of the interaction of the N-termini of various SUMO proteins with the respective SUMO cores and SIM-binding grooves, related to Figure 3.** (A) - (D) Displayed are various observables from MD simulations for the binding of the N-terminus to the core of wild-type SUMO1, SUMO2, Smo-1 and Smt3: (A) average binding area expressed as percent of the total area of the SUMO core, (B) average binding time expressed as percent of the total simulation time, (C) average binding area expressed in nm2 and (D) average binding area of the N-terminus not with the core, but with the SIM-binding groove expressed in $nm^2$. For each panel, data points represent the average along single 1 μs trajectories, bars are the average of 8 trajectories and errors bars are given as the standard errors over these 8 values; significance levels: * $p \leq 0.05$; ** $p \leq 0.01$; *** $p \leq 0.001$. (E) Distribution of binding/unbinding events for various SUMO proteins. Each row represents an analysis of binding/unbinding events of the N-terminus and the SUMO core for wild-type SUMO1, SUMO2, Smo-1 and Smt3, respectively. In each row, the first and third panel show the distribution of binding and unbinding events by residue, respectively, which were quantified by using the time evolution of binding areas shown in Figure S5C. Kinetic rate constants for binding and unbinding processes (second and fourth column in each row) were obtained by fitting a two-state model ($y = A^*(1 - \exp(-kx))$, red line) to cumulative histograms of binding/unbinding events over time (blue circles).

**Figure S6**

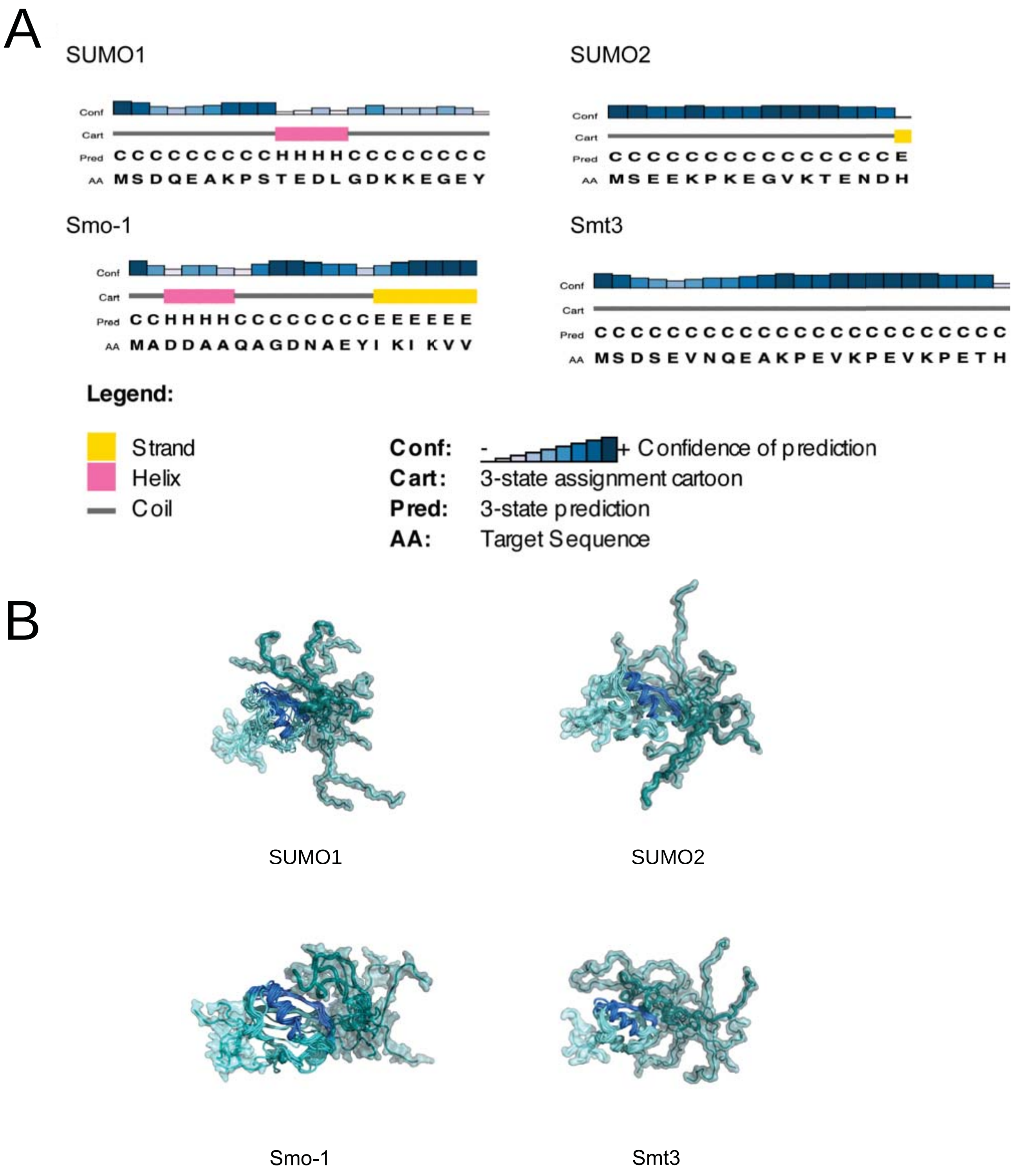
A
SUMO1
Conf
Cart
Pred CCCCCCCCCHHHHCCCCCCCC
AA MSDQEAKPSTEDLGDKKEGEY
SUMO2
Conf
Cart
Pred CCCCCCCCCCCCCCCE
AA MSEEKPKEGVKTENDH
Smo-1
Conf
Cart
Pred CCHHHHCCCCCCCCEEEEEE
AA MADDAAQAGDNAEYI KI KVV
Smt3
Conf
Cart
Pred CCCCCCCCCCCCCCCCCCCCCCC
AA MSDSEVNQEAKPEVKPEVKPETH
Legend:
Strand
Helix
Coil
Conf: - + Confidence of prediction
Cart: 3-state assignment cartoon
Pred: 3-state prediction
AA: Target Sequence
B
SUMO1
SUMO2
Smo-1
Smt3

**Figure S6 - continued**

C

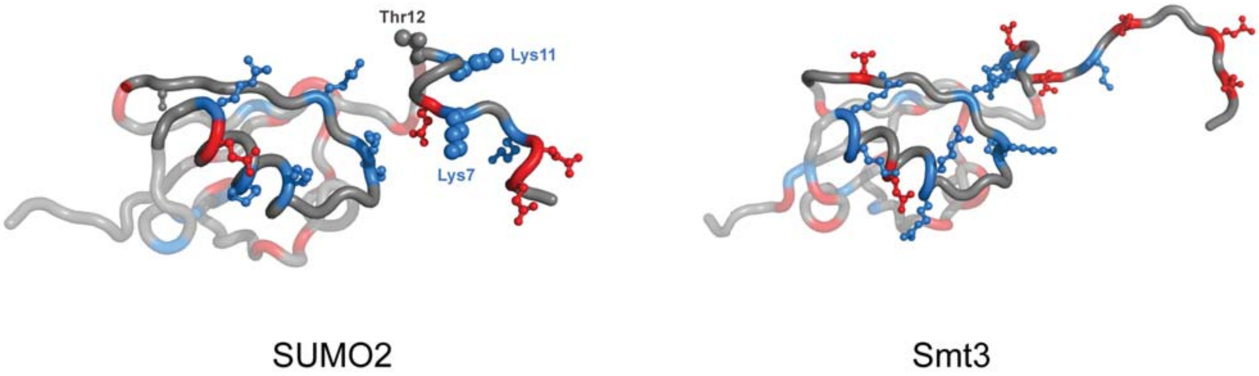

**Figure S6 Structural information about various SUMO proteins, related to Figure 3 and 4**

(A) Secondary structure prediction for the N-terminus of indicated SUMO protein with PSIPRED (Jones, 1999) reveals that they are predicted to be disordered regions.

(B) The N-termini of SUMO proteins behave as “ligand clouds” howering over the SUMO1 (upper left), SUMO2 (upper right), Smo-1 (lower left) and Smt3 (lower right) cores, respectively. Representative conformations of the N-termini for each respective protein are shown in turquoise, the SIM-binding groove is in blue and the rest of the protein in cyan.

(C) Highly mixed distribution of charged residues in SUMO proteins. Highly populated structural models clustered from MD simulations of human SUMO2 (left) and yeast Smt3 (right) are shown. The SIM binding groove, encompassing β2-strand and α-1 helix, as well as the N-terminus of each structure were shown in solid coloring, while the rest of the molecules is transparent. The positively and negatively charged residues were colored in blue and red, respectively, with their side chains represented as balls and sticks. The residues related with post-translational modifications in human SUMO2 are marked and shown as larger ball-and-stick models.

**Figure S7**

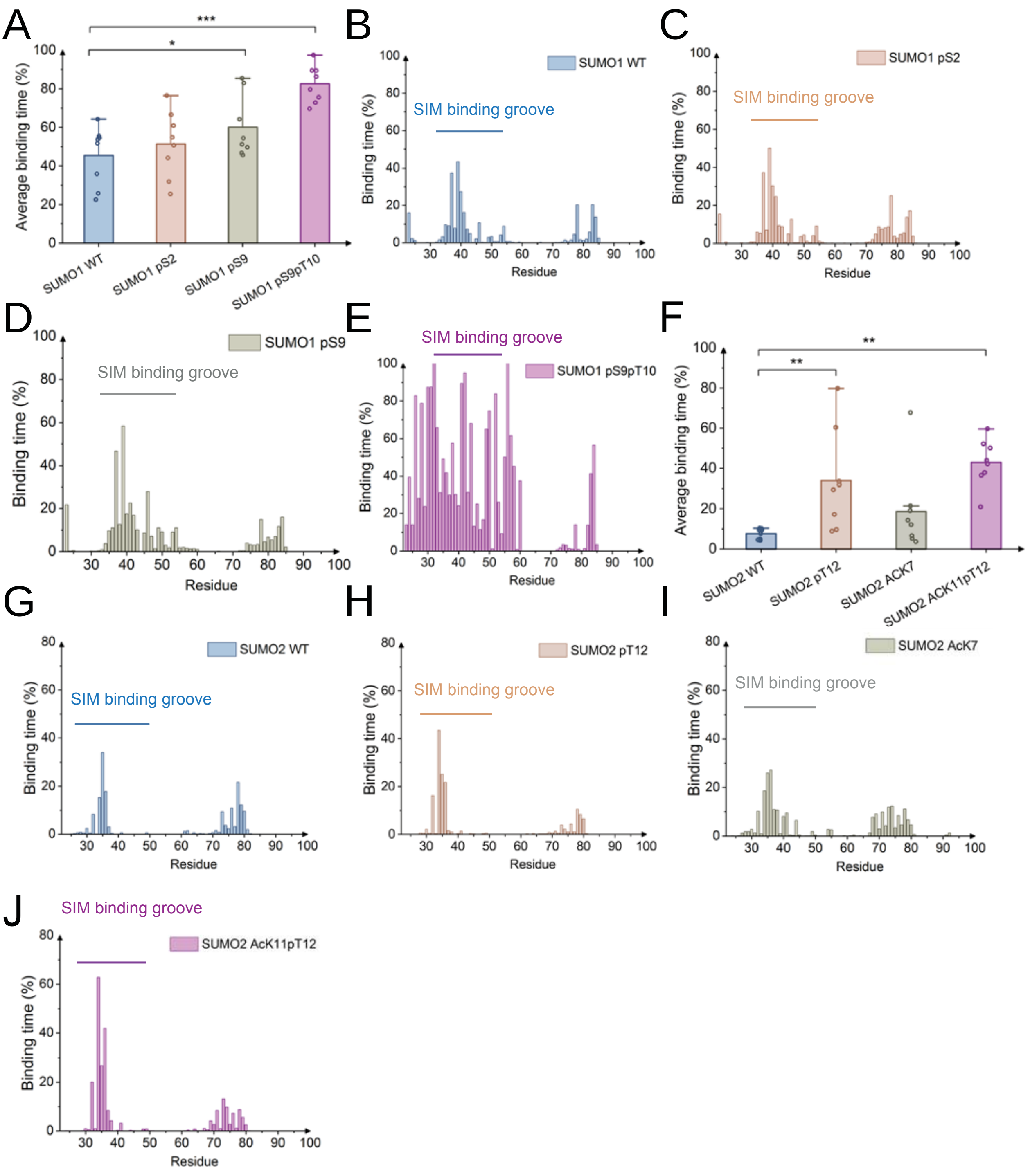
A
***
*
Average binding time (%)
SUMO1 WT
SUMO1 pS2
SUMO1 pS9
SUMO1 pS9pT10
B
SUMO1 WT
SIM binding groove
Binding time (%)
Residue
C
SUMO1 pS2
SIM binding groove
Binding time (%)
Residue
D
SUMO1 pS9
SIM binding groove
Binding time (%)
Residue
E
SIM binding groove
SUMO1 pS9pT10
Binding time (%)
Residue
F
**
**
Average binding time (%)
SUMO2 WT
SUMO2 pT12
SUMO2 ACK7
SUMO2 ACK11pT12
G
SUMO2 WT
SIM binding groove
Binding time (%)
Residue
H
SUMO2 pT12
SIM binding groove
Binding time (%)
Residue
I
SUMO2 AcK7
SIM binding groove
Binding time (%)
Residue
J
SIM binding groove
SUMO2 AcK11pT12
Binding time (%)
Residue

**Figure S7 Post-translation modifications of the intrinsically disordered region of SUMO proteins serve as a regulable cis-inhibitor for SIM-dependent interactions, related to Figure 6**

(A) Average binding time of the SUMO N-terminus to the SIM-binding groove for SUMO1 wt and the pS2, pS9 and pS9pT10 post-translationally modified SUMO1 variants. (B) - (E) Average binding time of the N-terminus of SUMO1 wt, pS2, pS9 and pS9pT10 to individual residues of the SUMO core. (F) Average binding time of the SUMO N-terminus to the SIM-binding groove for SUMO2 wt and the pT12, AcK7 and AcK11pT12 post-translationally modified SUMO2 variants. (H) - (J) Average binding time of the N-terminus of SUMO2 wt, pT12, AcK7 and AcK11pT12 to individual residues of the SUMO core. Data points represent the average along single 1 µs trajectories, columns are the average of 8 trajectories and errors bars are given as the standard errors of the mean of the 8 averages; significance levels: * $p \leq 0.05$; ** $p \leq 0.01$; *** $p \leq 0.001$. The panels shown in Fig. S7A, S7B, S7D and S7F - S7H are identical to those in Fig. 6A - 6F, respectively, and are displayed here again for comparative reasons.

**Table S2. Overview of worm strains incuding their genotypes used in this study, related to Figure 7.**

| **Strain** | **Genotype** | **Reference** |
|---|---|---|
| AH6024 | *smo-1(zh156) I* | this study |
| WS4116 | *bcIs39 [lim-7p::ced-1::GFP + lin-15(+)] V* | CGC*, Zhou et al., 2001 |
| AH6119 | *smo-1(zh156) I; bcIs39 [lim-7p::ced-1::GFP + lin-15(+)] V* | this study |
| AH5891 | *cep-1(gk138) I; bcIs39[Plim-7::CED-1::GFP] V* | Hajnal Lab |
| AH6143 | *smo-1(zh156) I; cep-1(gk138) I; bcIs39[Plim-7::CED-1::GFP] V* | this study |
| VC172 | *cep-1(gk138)* | CGC* |

* *C.elegans* Genetics Center (Minneapolis, MN)

**Figure S8**

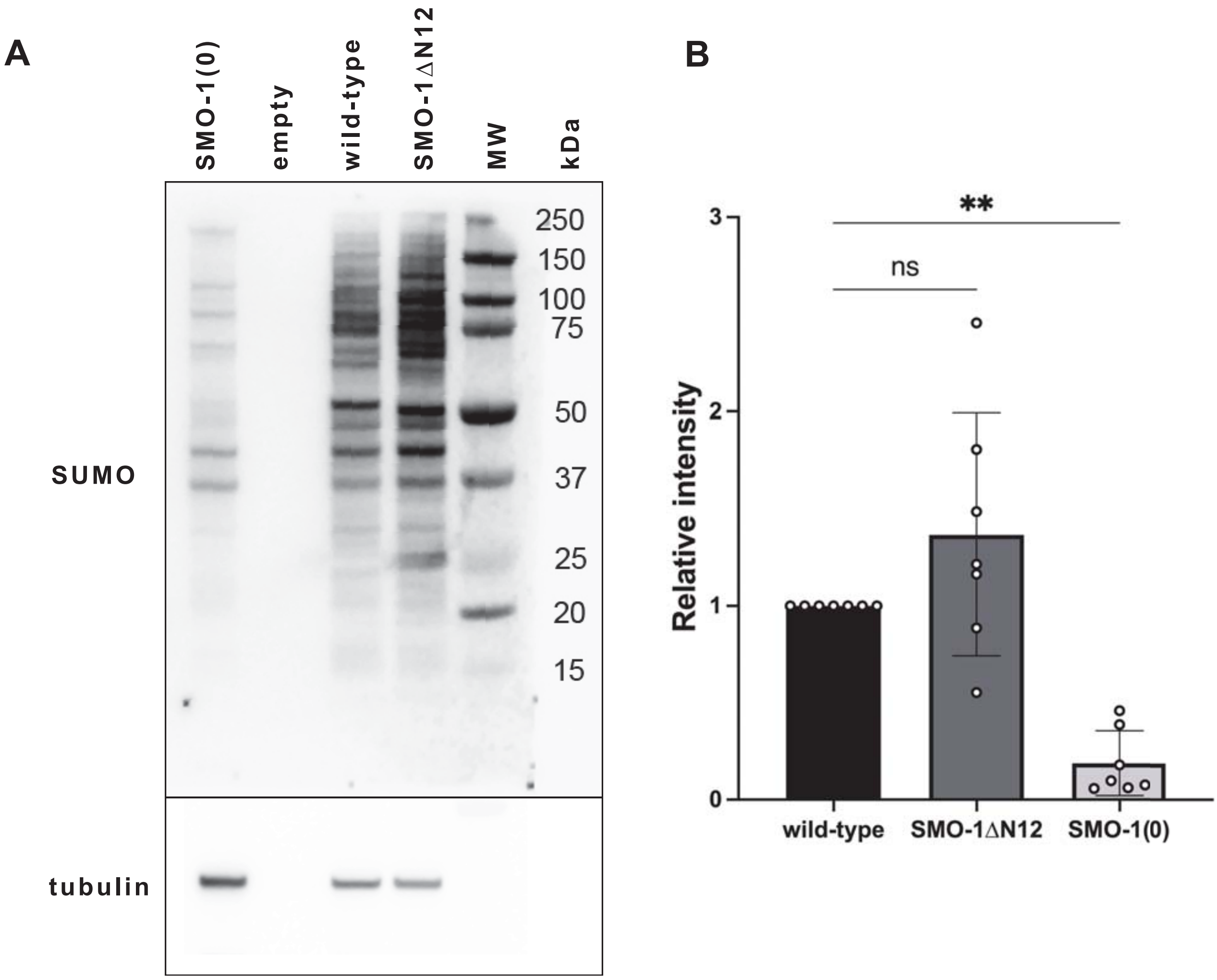
A
SMO-1(0)
empty
wild-type
SMO-1ΔN12
MW
kDa
250
150
100
75
50
37
25
20
15
SUMO
tubulin
B
**
ns
Relative intensity
3
2
1
0
wild-type
SMO-1ΔN12
SMO-1(0)

**Figure S8 SUMO levels in SMO-1ΔN12 animals compared to wild-type and SUMO deletion mutants**

Western blot of whole-animal extracts from one day-old adults probed with anti-SUMO (top) and anti-tubulin antibodies as loading control (bottom). SMO-1(0) refers to extracts from homozygous *smo-1(ok359)* SUMO null mutants and SMO-1ΔN12 to *smo-1(zh156)* extracts (A). Quantification of SUMO levels in 7 different samples from three biological replicates (B). Signal intensities were first normalized to the tubulin loading controls and then to the wild-type signal on each blot. Error bars indicate the standard deviations. Statistical significance was determined by one-way ANOVA with Dunnett's correction for multiple comparisons. ** indicates $p<0.01$ and n.s. $p>0.05$.
